# The microcephaly gene *ASPM* is required for the timely generation of human outer-radial glia progenitors by controlling mitotic spindle orientation

**DOI:** 10.1101/2023.09.25.559314

**Authors:** Anja van Benthem, Ridha Limame, Matteo Piumatti, Maurine Zaratin, Emir Erkol, Daisuke H. Tanaka, Adèle Herpoel, Angéline Bilheu, Harmen van Benthem, Laura Nebreda Rodriguez, Isabelle Pirson, Kathelijn Keymolen, Suresh Poovathingal, Martyna Ditkowska, Julie Désir, Sandrine Passemard, Marc Abramowicz, Catherine Ledent, Pierre Vanderhaeghen

**Affiliations:** Université Libre de Bruxelles (U.L.B.), Institut de Recherches en Biologie Humaine et Moléculaire (IRIBHM), and ULB Neuroscience Institute (UNI), 1070 Brussels, Belgium; VIB-KU Leuven Center for Brain & Disease Research, 3000 Leuven, Belgium; Department of Neurosciences, Leuven Brain Institute, KU Leuven, 3000 Leuven, Belgium; Centre for Medical Genetics, Reproduction and Genetics, Reproduction Genetics and Regenerative Medicine, Vrije Universiteit Brussel (VUB), UZ Brussel, Laarbeeklaan 101, 1090 Brussel; Institute of Pathology and Genetics (IPG), 25 avenue Georges Lemaître, 6041 Gosselies, Belgium; Université Paris Cité, Inserm UMR 1141, NeuroDiderot, F-75019 Paris, France; Service de Neurologie Pédiatrique, DMU INOV-RDB, APHP, Hôpital Robert Debré, F-75019 Paris, France; Cellistic, 1435 Mont-Saint-Guibert, Belgium; Altertox Academy, 1050 Brussels, Belgium; University of Geneva, Dept Genetics and Development, Faculty of Medicine, Rue Michel Servet 1, 1211 Geneva, Switzerland

## Abstract

Abnormal spindle-like, microcephaly-associated (*ASPM*) is the most commonly mutated gene in primary microcephaly (MCPH), characterized by reduced brain size and intellectual deficiency. The mechanisms underlying MCPH have remained unclear, as *ASPM* disruption leads to distinct phenotypes depending on the species studied. Here, we studied the impact of *ASPM* pathogenic mutations on human corticogenesis in organoid models. We found that at earliest stages of neurogenesis, *ASPM* mutant cortical progenitors, located at the apical surface of the neuroepithelium, display transient mitotic spindle randomization. The mutant progenitors then delaminate basally to adopt precociously the characteristics of outer radial glia cells (oRGC), a population of progenitors selectively amplified in human corticogenesis. Subsequently, cortical progenitors are depleted through decreased amplification and increased apoptosis. Thus, *ASPM* regulates the timely generation of oRGC by controlling mitotic spindle orientation, shedding light on how species-specific features of neurogenesis may confer vulnerability to neurodevelopmental diseases.

## Introduction

During recent hominin evolution, the cerebral cortex has undergone a rapid and considerable increase in size and complexity, with significant impact on the acquisition of higher functions in the human species ^1,2^. The increase in cortical size and neuronal number has been linked to evolutionary changes in the processes of generation of cortical neurons, or neurogenesis ^1,3,4^. Indeed, while cortical neurogenesis is well conserved among mammals, a number of divergent features have been identified during human cortical neurogenesis, in particular at the level of cortical stem and progenitor cells ^1,3^. Cortical progenitors constitute a diverse set of cells located first in the proliferative zones lining the lateral ventricles of the dorsal telencephalon, or ventricular zone (VZ). The first neural progenitors found in the cortical anlage are neuropithelial cells (NEC). NEC typically amplify by symmetric divisions and then convert into ventricular radial glial cells (vRGC), which divide and differentiate in the VZ, mostly through asymmetric divisions, and constitute the major subtype of neurogenic cortical progenitors throughout mammals ^5,6^. vRGC can convert directly into neurons, or expand further as intermediate progenitor (IP) cells, which divide more basally in the so-called subventricular zone (SVZ), thus further increasing the final neuronal output ^7,8^. vRGC can also convert into outer radial glial cells (oRGC), a subpopulation of neurogenic neural stem cells located further basally in the cortical anlage, in a special niche called the outer-subventricular zone (OSVZ). Importantly, this latter process is quantitatively and qualitatively divergent among mammals ^3,9^. While oRGC are found in very small numbers in rodents ^10,11^, they are much more numerous in carnivores and primates, and even more so in the human^12–14^. Human oRGC are characterized by highly enhanced amplification capacities^15^, which is thought to contribute importantly to increased human cortical size^12,15^. The generation of oRGC is a tightly controlled timely process ^16^, as it only occurs at later stages of cortical neurogenesis, thus contributing mostly to late-born upper layer neuron generation ^17^.

Thus, species-specific features of human corticogenesis have had a significant impact on human brain evolutionary expansion, but they could also have important implications for brain diseases, in which human-specific developmental mechanisms may be impaired. Among such diseases, non-syndromic primary microcephaly (MCPH) is a rare autosomal recessive condition characterized by a severely decreased brain size at birth ^18,19^. It is clinically described as a reduction of occipitofrontal head circumference of at least two standard deviations below the mean for age, gender and ethnicity ^20^, and in particular a smaller cerebral cortex ^18^. MCPH can be caused by mutations in at least 30 genes ^19^ (https://www.omim.org/entry/620183?search=MCPH30&highlight=mcph30#1). *ASPM* is the most frequently mutated gene in MCPH patients ^21^. It encodes a pericentrosomal protein that displays several functional modules including a large IQ-rich domain (Figure 1A), and that is highly enriched at the level of the mitotic spindle in most cell types ^22,23^. *ASPM* gene orthologues are found throughout the animal kingdom, but *ASPM* sequence was found to display signs of accelerated evolution and/or positive selection, as well as an increase in the size of its IQ-rich domain in vertebrates and primates ^24,25^. In *C. elegans* and *Drosophila*, ASPM orthologues are required to control cytokinesis and mitotic spindle organization in several cell types ^26–30^. Acute knock-down of *aspm* in zebrafish results in mitotic arrest and modest brain size reduction ^31,32^, while in mouse embryonic cortex *ASPM* knock-down leads to precocious neuronal generation, together with misregulation of the angle of mitotic spindle and cell division ^33,34^. However genetic models of mice displaying similar mutations as those found in patients display only very mild microcephaly ^35–37^, and no detectable or weak phenotypes related to mitotic spindle and/or neurogenesis ^35–38^. These data suggest that mice may display lower sensitivity to disrupted ASPM function, compared with the more dramatic effects seen in the human, suggesting some degree of species-specific sensitivity. Consistent with this hypothesis, *ASPM* gene knock-out leads to microcephaly in the ferret ^39^, a species in which cortical neurogenesis is more closely related to the human, including the abundant presence of oRGC^14^. In addition to strong cortical volume reduction, *Aspm* knock-out in the ferret was associated with a premature delamination of cortical progenitors to the OSVZ ^39^.

**Figure 1.**
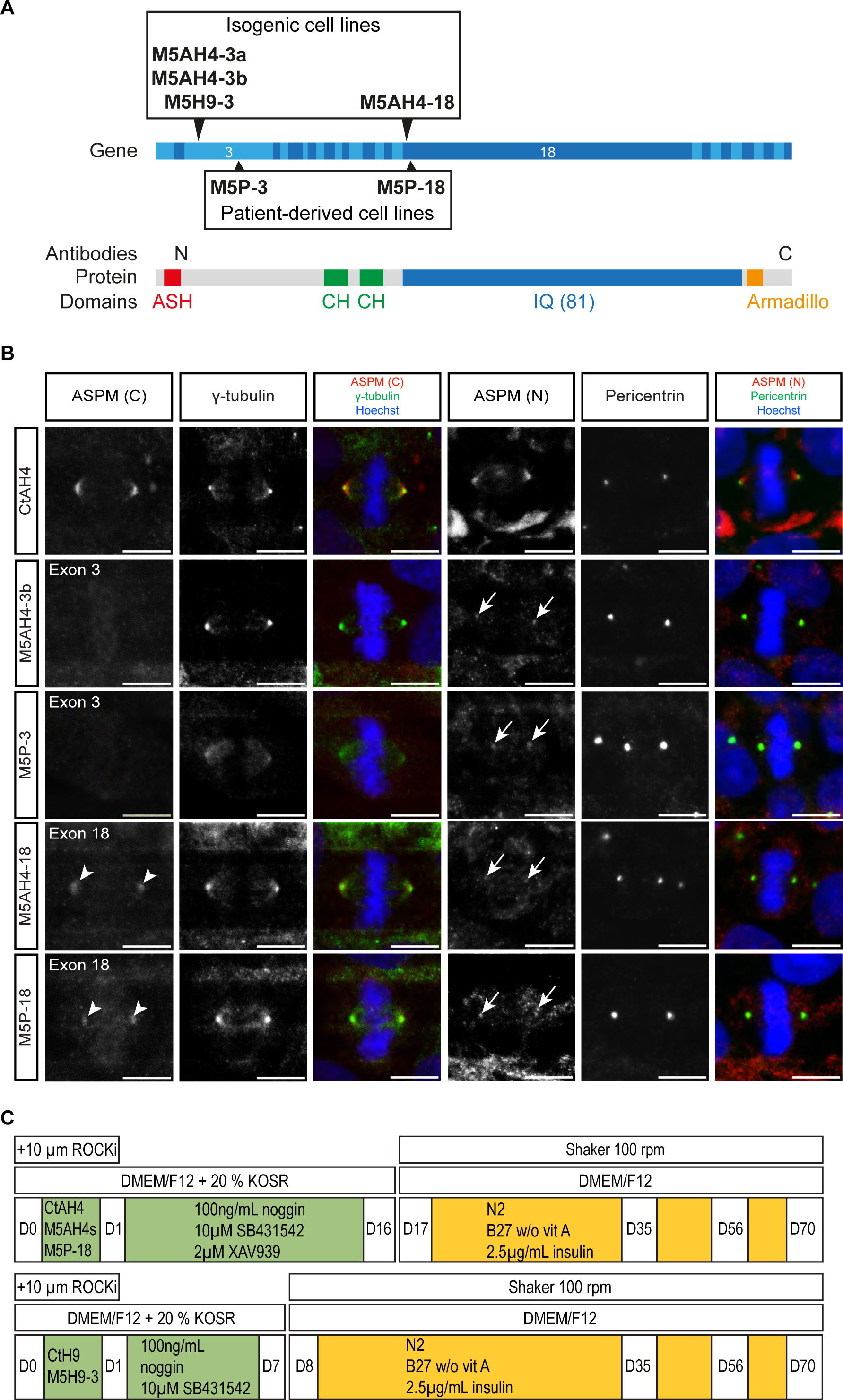
*ASPM*-mutated hiPSC/hES cell lines and cortical organoid differentiation system. (A) Structure of the 28 human *ASPM* exons (top) encoding the ASPM protein (bottom). Position of patient’s *ASPM* mutations (M5P-3 and M5P-18, box below *ASPM* gene, see also Table S2). Position of CRISPR/Cas9-mediated introduction of homozygous mutations by insertion of a T in exon 3 (c.699insT) or exon 18 (c.4199insT) in the control hiPSC lines CtAH4 and the control hESC line CtH9 are shown as pairs of black arrowheads (box above *ASPM* gene, see also Table S1). Domains of the 3477 amino acid comprising ASPM protein are shown: an N-terminal ASH (ASPM, SPD-2, Hydin) domain with putative microtubule-binding function (red), two calponin homology (CH) domains with actin-binding funtion (green), 81 calmodulin-binding IQ domains (blue) and a C-terminal armadillo repeat-like domain (orange) ^23,43^. The epitopes of the ASPM antibodies directed against the N- and C-terminus are depicted on top of the ASPM protein structure (shown as N and C, respectively). (B) Subcellular localization of ASPM protein in mitotic hiPSCs of a control individual (CtAH4), its isogenic hiPSCs with a mutation in exon 3 (M5AH4-3b) and exon 18 (M5AH4- 18) and patient iPSCs with similar mutations in exon 3 (M5P-3) and exon 18 (M5P-18). In control hiPSCs, ASPM immunostaining (merged image: red) is surrounding and partially overlapping with the centrosomal γ-tubulin or Pericentrin staining (merged image: green) at the spindle poles. Merged images additionally show Hoechst staining (blue). Immunoreactivity of the C-terminal ASPM antibody is weaker in M5AH4-18 hiPSCs than in their isogenic control, whereas no signal is found at the spindle poles of M5AH4-3b hiPSCs, as shown in single channel images (grey). The N-terminal ASPM antibody shows a weaker signal at the spindle poles in *ASPM*-mutated compared with control hiPSCs. Scale bars, 10 µm. See also Figures S4B and S4C. (C) Schematic drawing of the cortical organoid differentiation system *in vitro* for the isogenic AH4 hiPSC lines and the patient-derived hiPSC line M5P-18 (top). For the isogenic H9 hESC lines (bottom) the neural induction phase is shortened to only eight instead of 16 days with simultaneous increase of the concentration of Wnt inhibitor XAV939 to 5 µM instead of 2 µM. See also Figure S1.

*ASPM* may thus display species-specific functions in species with increased cortical size. However, it has remained unclear how *human* corticogenesis is altered by *ASPM* mutations. *ASPM* mutations were examined in neural organoids generated from patient-derived iPSC and were found to cause strong defects in early organoid growth ^40^. But these phenotypes and the underlying mechanisms have remained difficult to interpret, as the regional identity of the organoids generated was unclear, and no isogenic control was examined.

Here we studied the impact of pathogenic mutations of *ASPM* in human corticogenesis in cortical organoid models. Using human pluripotent stem cells (PSC) bearing several types of pathogenic mutations of *ASPM* and their respective isogenic controls, we found that in all cases the mutated forms still encode a protein that is localized at the mitotic spindle, but consistently lacks the IQ-rich domain. Mutant cortical NEC and vRGC displayed a precocious randomization in mitotic spindle and cell division orientation, followed by the early delamination of these progenitors from the VZ to the basal compartment, and acquisition of an oRGC-like cellular phenotype. The generation of oRGC ahead of time was accompanied by increased apoptosis and depletion of vRGC. Our findings reveal that mutations in *ASPM* result in defective corticogenesis through alterations in the timing of critical cell transitions in human cortical progenitors, providing a mechanistic link between *ASPM* function in the developing human neocortex and MCPH pathogenesis.

## RESULTS

### *ASPM* pathogenic mutations lead to truncated forms of ASPM proteins

To study the impact of MCPH-related *ASPM* mutations in different genetic backgrounds and using isogenic controls, we generated iPSC (CtAH4)^41^ and ESC (H9) lines displaying homozygous ASPM mutations using CRISPR/Cas9 gene editing. We thus targeted ASPM exon 3 and exon 18, leading to frameshifts closely similar to those found in MCPH patients (Figures 1A and S1A, Tables S1 and S2). In addition, we established iPSC lines reprogrammed from dermal fibroblasts obtained from MCPH patients carrying mutations in the *ASPM* gene in the same exons (M5P-3 and M5P-18) (Figures 1A, S1A-E, and Table S2). *ASPM* genotype and chromosomal integrity were confirmed in all cell lines by sequencing and karyotyping, and pluripotency maintenance was confirmed using *ad hoc* markers and embryoid body formation (Figures S1A-E, and data not shown).

We next characterized the expression of ASPM protein in control and mutant PSC. We used two antibodies, directed against the C- or the N-terminal end of ASPM (Figures 1B and S1F). This revealed expression of the ASPM protein in control PSC at the level of the mitotic spindle poles, as described previously ^23,42^. The ASPM signal surrounded, and partially overlapped with, the γ-tubulin and Pericentrin stainings, which label centrosomes (Figures 1B and S1F). We next examined *ASPM*-mutated PSC. Interestingly, this revealed that ASPM protein immunoreactivity was still detectable at the spindle pole, at variable levels depending on the exon targeted by the mutation. In PSC with exon 3 *ASPM* mutations (M5AH4-3a and -3b, M5H9-3, and M5P-3), we detected no immunoreactivity with the C-terminal antibody, as expected, but adid detect a weak immunoreactivity with the N-terminal antibody, localized at the spindle pole level (Figures 1B and S1F, indicated by arrows). On the other hand, in PSC with exon 18 *ASPM* mutations (M5AH4-18, M5P-18), we detected control-like immunoreactivity for the ASPM protein using both antibodies (Figures 1B, C-terminal signal indicated by arrowheads, and S1F). This result can be explained by differential exon 18 usage in the exon 18-mutant cells, in which the exon 18 would be skipped preferentially when mutated ^43^. Overall these results suggest that distinct pathogenic mutations of *ASPM*, in either exon 3 or 18, do not lead to complete loss of the protein, but rather to truncated forms, all of which lack the exon 18-encoded IQ domain, but that can still localize to the mitotic spindle pole.

## Generation and validation of cortical organoids

To study the impact of *ASPM* mutations on cortical neurogenesis *in vitro*, we established a cortical organoid differentiation protocol (Figure 1C), starting from whole PSC colonies and based on dual SMAD inhibition of TGFβ and BMP by SB431542 and Noggin respectively, combined with WNT inhibition by XAV939 during the neural induction phase ^44,45^. Importantly, the duration and concentration of XAV939 was adjusted depending on the parental cell line (AH4 or H9) to maximize the outcome of cortical fate (see Methods). The colonies self-assembled in floating aggregates in non-adherent culture plates and developed further into translucent, radially organized VZ-like structures (Figure S2). We first validated the protocol on the two control AH4 and H9 lines (Figures S2 and S4A). PSC differentiated into organoids containing multiple cell-dense ventricular zone-like (VZ-like) structures (Figures S2A–B). Most of the cells of these structures co-expressed SOX2, PAX6, FOXG1 and EMX1, indicative of dorsal telencephalon or cortical identity, although a smaller number of VZ-like structures were found to be negative for FOXG1/EMX1, suggestive of other regional identity (Figure S2). The VZ-like structures displayed apicobasal polarity, revealed by the apical protein complex marker ZO-1, and apical enrichment of mitotic phospho-histone 3 (PH3)+ progenitor cells, suggestive of interkinetic nuclear migration as found *in vivo* (Figure S2B). Notably, in addition to the VZ-like structures with cortical identity, the organoids displayed a contingent of cells expressing OTX2, a marker for the choroid plexus (Figures 2A, S2A, and S3B) ^46^. Around day 35, TBR2+ (also known as EOMES) IPC could also be found more basally next to the VZ-like structures, thus mimicking an SVZ-like domain (Figure S2B). The progenitor domains were further surrounded by neurons expressing EMX1 and FOXG1, as well TBR1 and CTIP2 (also known as BCL11B), indicative of deep-layer neuron identity. At later stages (from day 70), an additional contingent of SATB2+ or Cux1+ neurons could be observed (Figures 2C–E, and S2B), indicative of upper layer neuron identity. The cortical organoids thus display a conserved temporal patterning of cortical neurogenesis, as expected ^41,47,48^. Notably however, the neurons expressing deep-layer and upper-layer neurons appeared to be mostly intermingled, indicating that this system could not faithfully recapitulate the upside down generation of cortical layers found *in vivo*.

**Figure 2.**
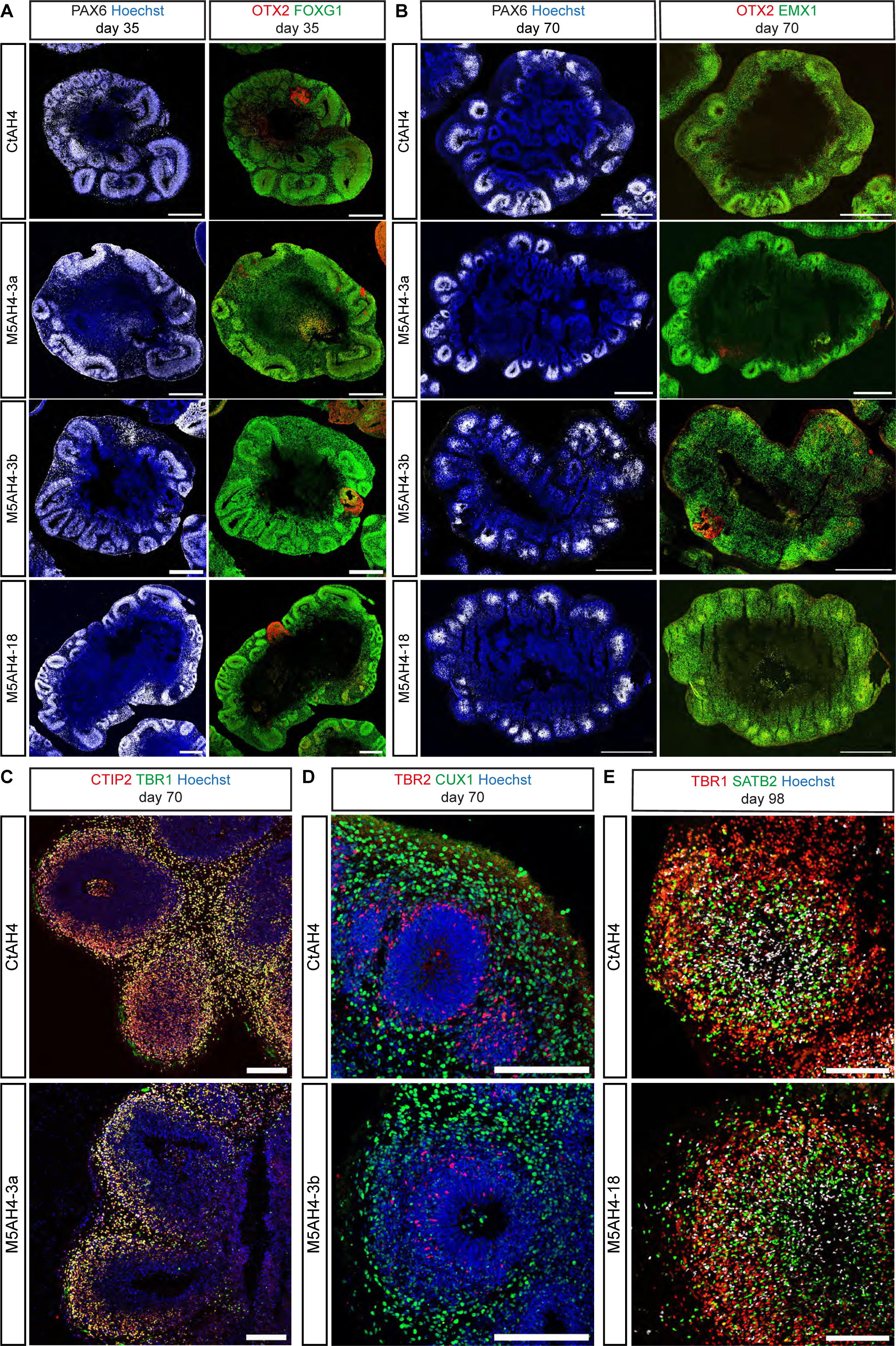
Efficient generation of cortical organoids from control and *ASPM*-mutated hPSCs. (A and B) Immunofluorescence staining showing that most of the VZ-like structures of organoids derived from control (CtAH4) and *ASPM*-deficient (M5AH4) hiPSCs are positive for the neural progenitor marker PAX6, the forebrain marker FOXG1 (A), and the dorsal telencephalon marker EMX1 (B). Some regions are positive for OTX2, a marker of choroid plexus and diencephalon, and are typically negative for FOXG1, EMX1 and PAX6. (C and D) Immunofluorescence staining showing cortical deep-layer neurons (TBR1+/CTIP2+) at day 70 in control and *ASPM*-mutated conditions (C) and superficial-layer neurons (CUX1+) (D) outside of the VZ-like structures, while the intermediate progenitor marker TBR2 is found basally adjacent to the VZ-like structure, as described earlier ^70^. (E) Immunofluorescence staining showing substantial loss of VZ-like cytoarchitectural organization in organoids by day 98, although their center is still mainly populated by Sox2+ progenitor cells. In control and *ASPM*-mutated conditions, many SOX2+ cells are also intermingled with the surrounding TBR1+ deep-layer and SATB2+ upper-layer neurons. Scale bars, 500 µm (A), 1 mm (B), 200 µm (C-E). See also Figures S2–S4.

We next tested the validity of the organoid model using single cell RNA sequencing (scRNAseq), focusing on day 35 organoids. Based on unsupervised clustering we identified 17 cell types according to marker gene enrichment ^45^ (Figures S3A and S3E). These contained clusters corresponding to progenitors and neurons expressing markers of the cortical excitatory neuron lineage (FOXG1+/ EMX1+), choroid plexus-like cells (Figures S3A–C), and a smaller contingent of other cell types that could be assigned to other brain regions (Figures S3A, S3C, and S3E), as previously described ^45^. Importantly, the cells of FOXG1+/ EMX1+ organoid clusters could be faithfully mapped with those of *in vivo* human fetal cortex at CS22 ^49^, further supporting recapitulation of *in vivo*-like gene expression in the FOXG1/EMX1+ cells (Figure S3D).

## Corticogenesis in *ASPM*-mutant and isogenic cortical organoids

We then used our organoid system to directly compare corticogenesis in control and *ASPM*-mutant cells (Figure 2, Figure S2 and S4A). Importantly, due to the observed regional heterogeneity during cortical organoid differentiation (Figures S2A and S3B), we focused our analyses exclusively on organoid domains where expression of dorsal telencephalic markers could be well documented (i.e. FOXG1- or EMX1-positive VZ-like structures and their immediate basal surroundings).

Organoid characterization (from day 35 to 70) was performed on three *ASPM*-mutant AH4 hiPSC lines (M5AH4-3a, M5AH4-3b, and M5AH4-18) and their isogenic control (CtAH4), one *ASPM*-mutant H9 cell line (M5H9-3) and its isogenic control (CtH9), and one patient derived cell line (M5P-18) (Figure 1A). Importantly, all the similarities and differences that we observed between control and *ASPM*-mutant cells were found to be consistent across genetic background and cell lines.

We first examined the expression of ASPM protein in control and mutant cortical organoids using the C-terminal antibody. For the N-terminal antibody we stained 2D- differentiated cortical progenitors^41^ because of poor compatibility with the required methanol fixation conditions. These confirmed our observations made in PSC, indicating that ASPM protein was still expressed at the level of the mitotic spindle, but as a truncated variant, in all mutant backgrounds (Figure S4B and S4C).

Next we examined the overall organization and cellular composition of the organoids, which appeared to be comparable in all cell lines irrespective of the *ASPM* mutation (Figures 2A–B and S4A). All cell lines enriched over time for cortical fates as shown by their expression of EMX1 and FOXG1 (Figures 2A–B) and generated TBR2+ intermediate progenitors as well as cortical deep-layer (TBR1, CTIP2), followed by upper-layer (CUX1, SATB2) neurons (Figures 2C–E). These data indicate that the global patterns of morphogenesis and neurogenesis appear to be normal in *ASPM* mutant organoids. This is in contrast with the data previously reported of severely disrupted growth of organoids following *ASPM* disruption ^40^, which could be linked to the lack of isogenic control and unclear cell identity in the organoids examined in this study.

## Premature randomization of mitotic spindle orientation in *ASPM*-deficient cortical progenitors at early developmental stages

ASPM was previously reported to be implicated in mitotic spindle orientation, with variable impact depending on the species and cells examined ^22,27,34,38^. We thus examined the mitotic spindle in dividing vRGC, located at the apical side of cortical (FOXG1+) VZ-like structures in isogenic control and *ASPM*-mutated cortical organoids (Figures 3 and S5). We focused on anaphase and early telophase, since nucleus and spindle can undergo wide oscillating movements in metaphase, but stop rotating in anaphase ^50,51^. At early stage (day 35), control AH4 and H9 cortical progenitors divided mostly (>95%) with a horizontally oriented spindle, parallel to the apical surface (Figures 3A–D, S5A), as typically found in mammalian cortex *in vivo* ^50,52–55^. At later stages, they gradually displayed randomized patterns, with an increased number of cells displaying vertical spindle pole orientation, thus leading to more and more cells displaying a horizontal plane of division (Figure 3A–D), consistent with previous findings in human fetal cortex ^52^. On the other hand, examination of the *ASPM*-mutated cells at day 35 revealed an early randomization in the orientation of the mitotic spindle, leading to a large increase in the number of spindle orientations with non-horizontal and even vertical cleavage angle divisions at this early stage (Figures 3A–D, and S5A). At later stages, the mitotic spindle pole angle remained randomized, so that control and mutant cells displayed similar ranges of mitotic spindle angle variability (Figure 3B–D, and S5A).

**Figure 3.**
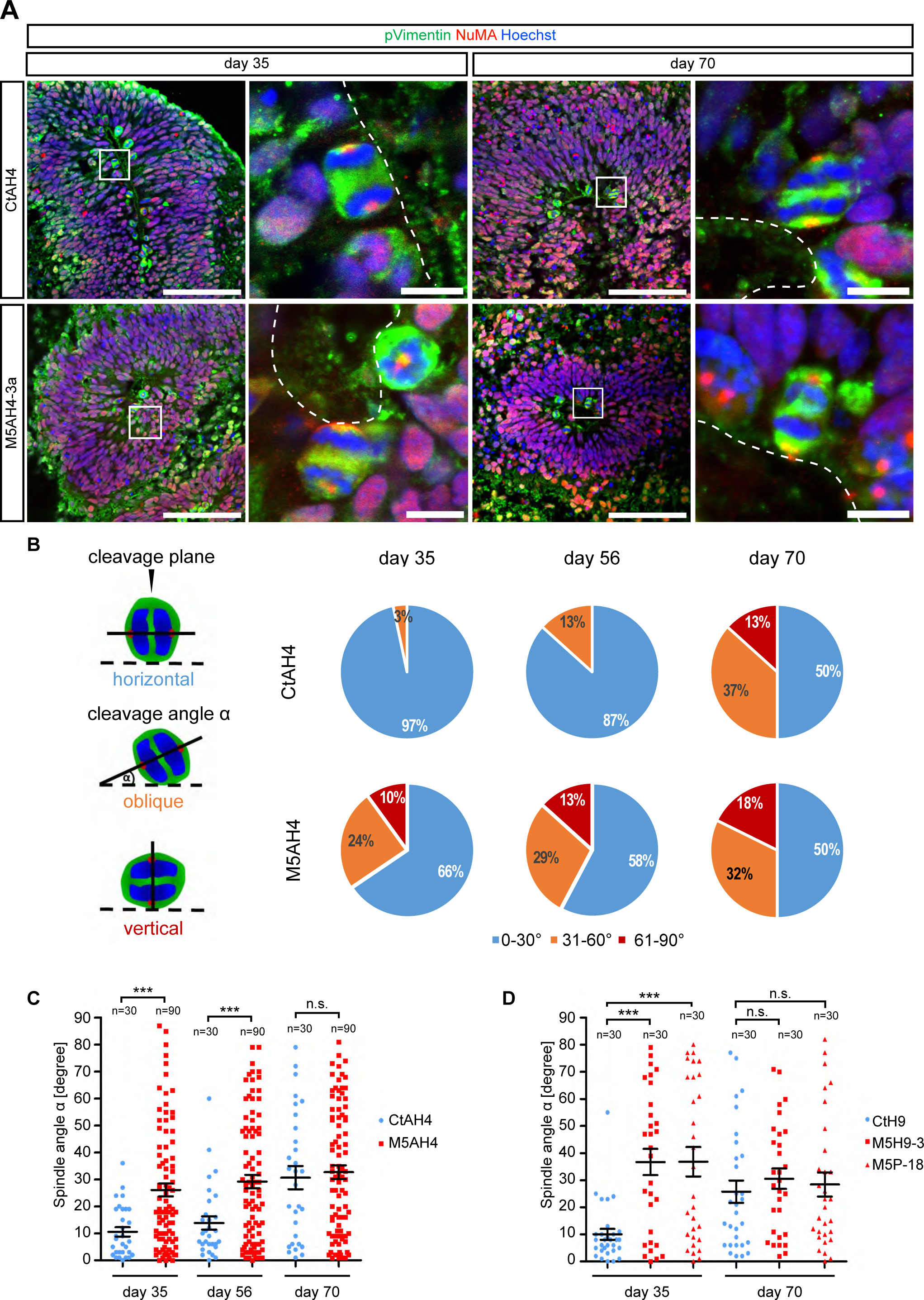
Premature alteration of spindle orientation in *ASPM*-deficient neural progenitors of cortical organoids. (A) Representative confocal images of mitotic neural progenitors in anaphase in organoids derived from isogenic control and *ASPM*-deficient AH4 hiPSCs at day 35 (left) and day 70 (right). Shown are overview immunofluorescence stainings for pVimentin (green) and NuMA (red) on the left with higher magnification of the boxed area in the right panel. Dashed lines indicate the apical surface. While most of the mitotic cells in the control condition CtAH4 divide in parallel to the apical surface, the number of cells with an oblique/vertically oriented cleavage angle is increased in *ASPM*-deficient cells (M5AH4-3a) at day 35. However, at day 70, the spindle orientation of control cells was found similarly altered to more oblique/vertical cleavage angles as in *ASPM*-mutated cells. (B, left) Schematic of dividing cells in relation to their apical surface (dashed line). The cleavage angle α is defined as spindle orientation and can be calculated computationally after 3D reconstruction of confocal images of the dividing cell. It reflects the orientation of a vector defined by the coordinates of the center of mass of each chromatin half of the mitotic cell in relation to the apical surface. (B, right) Pie diagrams of the proportion of dividing isogenic AH4 progenitor cells with horizontal (α = 0–30°), oblique (α = 31–60°) and vertical (α = 61–90°) cleavage angles in organoids at day 35, 56 and 70 analyzed by 3D reconstruction. The proportion of oblique and vertical cleavage angle orientations increases over time in both, control and *ASPM*-mutated cells; however, at day 35, it is already at an increased level in *ASPM*-mutated cell lines compared with the control cell line. (C) Scatter plot showing the quantification of cleavage angle orientation in control and *ASPM*-deficient AH4 neural progenitor cells analyzed by 3D reconstruction at day 35, 56 and 70 of organoid differentiation. One dot represents one cell. At day 35, most of the cortical progenitors of the control group divide horizontally (mean α = 10.57 ± 1.74 SEM), while *ASPM*-mutated progenitors show a premature increase in the distribution of oblique/vertical cleavage angles (mean α = 26.09 ± 2.38 SEM). At day 56, the cleavage angle is still predominantly horizontally oriented in control conditions (mean α = 13.83 ± 2.45 SEM), while significantly increased in *ASPM*-mutated (mean α = 29.24 ± 2.45 SEM) conditions. At day 70, the spindle orientation of the control cells (mean α = 30.67 ± 4.32 SEM) aligns with that of *ASPM*-mutated cells (mean α = 32.68 ± 2.50 SEM). Ten cells were analyzed per cell line and time point in three experiments using one control (CtAH4) and three isogenic *ASPM*-mutated cell lines (M5AH4-3a, -3b, -18). (D) Scatter plot showing the quantification of cleavage angle orientation in isogenic control and *ASPM*-deficient neural progenitor cells of H9 and patient-derived M5P-18 cells at day 35 and 70 of organoid differentiation. One dot represents one cell. At day 35, most of the cortical progenitors of the control group divide with a horizontal cleavage angle (CtH9: mean α = 10.03 ± 2.00 SEM), while *ASPM*-mutated progenitors show a premature increase in cells with an oblique/vertical cleavage angle (M5H9-3: mean α = 36.70 ± 4.85 SEM; M5P-18: mean α = 36.80 ± 5.39 SEM). At day 70, the number of oblique/vertical cleavage angles in control cells (CtH9: mean α = 25.77 ± 4.14 SEM) reaches a similar level as in *ASPM*-mutated cells (M5H9-3: mean α = 30.53 ± 3.77 SEM; M5P-18: mean α = 26.8 ± 4.98 SEM). Ten cells analyzed per cell line and time point in three experiments. All analyzed cells in were located in VZ-like structures with cortical identity as ensured by FOXG1 staining on consecutive sections (not shown). Mean ± SEM; Student’s *t*-test. Scale bars, 10 µm. See also Figure S5.

These results indicate that *ASPM* mutant cells display a precocious randomization of the mitotic spindle pole orientation. To study these events at even earlier stages we turned to another 3D protocol using corticospheres generated in Matrigel droplets (3DM) (Figure S5B). In this system, human PSC differentiated into neuroepithelial “corticospheres” within a few days, with a central apical lumen reminiscent of a ventricle and displayed robust morphological features of the developing neuroepithelium (Figure S5C) with typical apicobasal polarity (Figure S5D). The centrosomal markers Pericentrin and ψ-tubulin were found at the spindle poles of mitotic PH3-positive cells dividing close to the apical surface (Figure S5D). Furthermore, whilst cells in M-phase were consistently found at the apical side of the spheres, cells in S-phase (labeled by a short pulse of BrdU) were located at their basal surface (Figure S5E), indicating stereotyped interkinetic nuclear migration^56^. Over time, the size of the 3DM corticospheres increased until day 21, when the first β-III- tubulin/TBR1-positive neurons could be detected (Figure S5F) ^57^. Using this system mimicking the earliest stages of NE amplification, we directly compared control and *ASPM*-mutant cells. The overall organization and cellular composition of the 3DM spheres appeared comparable during the first three weeks (Figure S5C). We next examined the spindle orientation (Figures S5G–I). While the control lines displayed a robust horizontal cleavage plane at all examined stages, the *ASPM*-mutated neural progenitors revealed a randomization in the orientation of the mitotic spindle leading to mostly non-horizontal cleavage angle divisions (Figure S5 G-I).

Taken together, our data indicate that early human NE cells display a robust horizontal cleavage plane orientation, which is disrupted in the *ASPM*-mutated NEC, leading to premature randomization of the cleavage plane at the early stages of human corticogenesis.

## Precocious delamination of SOX2+ cortical progenitors from the VZ in *ASPM*- mutated organoids

We next examined in more detail the cellular distribution and localization of the cells in the organoids following randomization of the mitotic spindle pole. Remarkably, this revealed an early appearance of SOX2+ neural progenitor cells in basal compartments of mutant organoids, suggesting delamination of vRGC (Figure 4A, Figure S6A,B). Further examination and quantification over time revealed excessive and precocious delamination of SOX2+ vRGC, mimicking the generation of oRGC normally occuring at later stages (Figures 4A-C and S6A,B). At later stages (day 70), this also led to disruption of the cytoarchitecture of the VZ-like, with less well-defined boundaries between VZ and more basal compartments (Figure 4A, dashed lines).

**Figure 4.**
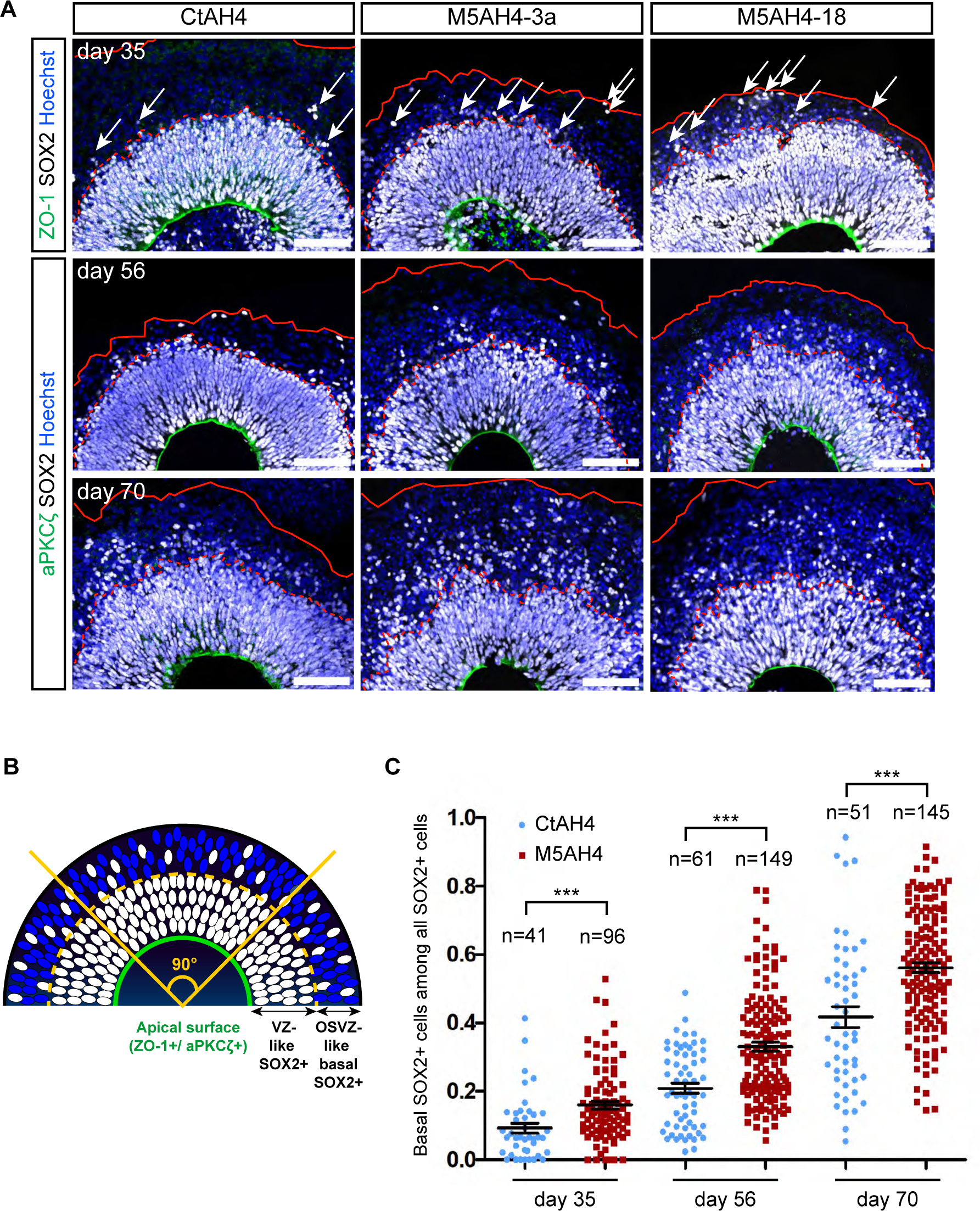
Precocious delamination of SOX2+ cortical progenitors from the VZ in *ASPM*-mutated organoids. (A) Immunostaining showing SOX2+ neural progenitors (white) and the apical surface labeled by ZO-1 or aPKCσ (green) in the VZ-like zones of isogenic AH4 control and *ASPM*-deficient organoids at day 35, 56 and 70. Representative confocal images of one control and two isogenic *ASPM*-deficient AH4 cell lines, mutated in exon 3 and exon 18. The SOX2+ neural progenitor cells delaminate prematurely form the VZ in *ASPM*-mutated organoids at all three time points. All VZ-like structures were located in regions with cortical identity, as confirmed by FOXG1 staining on consecutive sections. Dashed red lines show the border between VZ and the basal compartment, continuous red lines demarcate the outer surface of the organoids. (B) Schematic drawing of a cell-dense VZ-like zone with SOX2+ progenitor cells (white) lined by the apical surface (green) and with a basally adjacent OSVZ-like zone containing (more loosely packed) neurons (blue) and scattered SOX2+ progenitors. To quantify the number of SOX2+ cells, an angle of 90° was selected on confocal images open to the outer border of the organoid (yellow line). (C) Scatter plot showing the quantification of basal SOX2+ cells delaminated from the VZ- like zone among the total amount of SOX2+ cells within selected 90° angles in isogenic AH4 organoids at day 35, 56, and 70. One control and three pooled isogenic *ASPM*- deficient AH4 cell lines from at least four experiments were analyzed. Day 35: *P* value = 0.0007; CtAH4: 41 VZs from 11 organoids from 4 experiments, 3,919 SOX2+ cells; M5AH4s: 96 VZs from 41 organoids from 5 experiments, 26,914 SOX2+ cells. Day 56: *P* value < 0.0001; CtAH4: 61 VZs from 14 organoids from 6 experiments, 10,688 SOX2+ cells; M5AH4s: 149 VZs from 32 organoids from 5 experiments, 30,924 SOX2+ cells. Day 70: *P* value < 0.0001; CtAH4: 51 VZs from 15 organoids from 6 experiments, 8,079 SOX2+ cells; M5AH4s: 145 VZs from 33 organoids from 5 experiments, 24,712 SOX2+ cells. See also Figures S6A–B.

We further explored the identity of the the aberrantly delaminated SOX2+. This revealed that many of the displaced SOX2-positive cells co-expressed the oRGC-markers TNC, LIFR, and HOPX (Figures 5A,B and, S6C)^15^.

**Figure 5.**
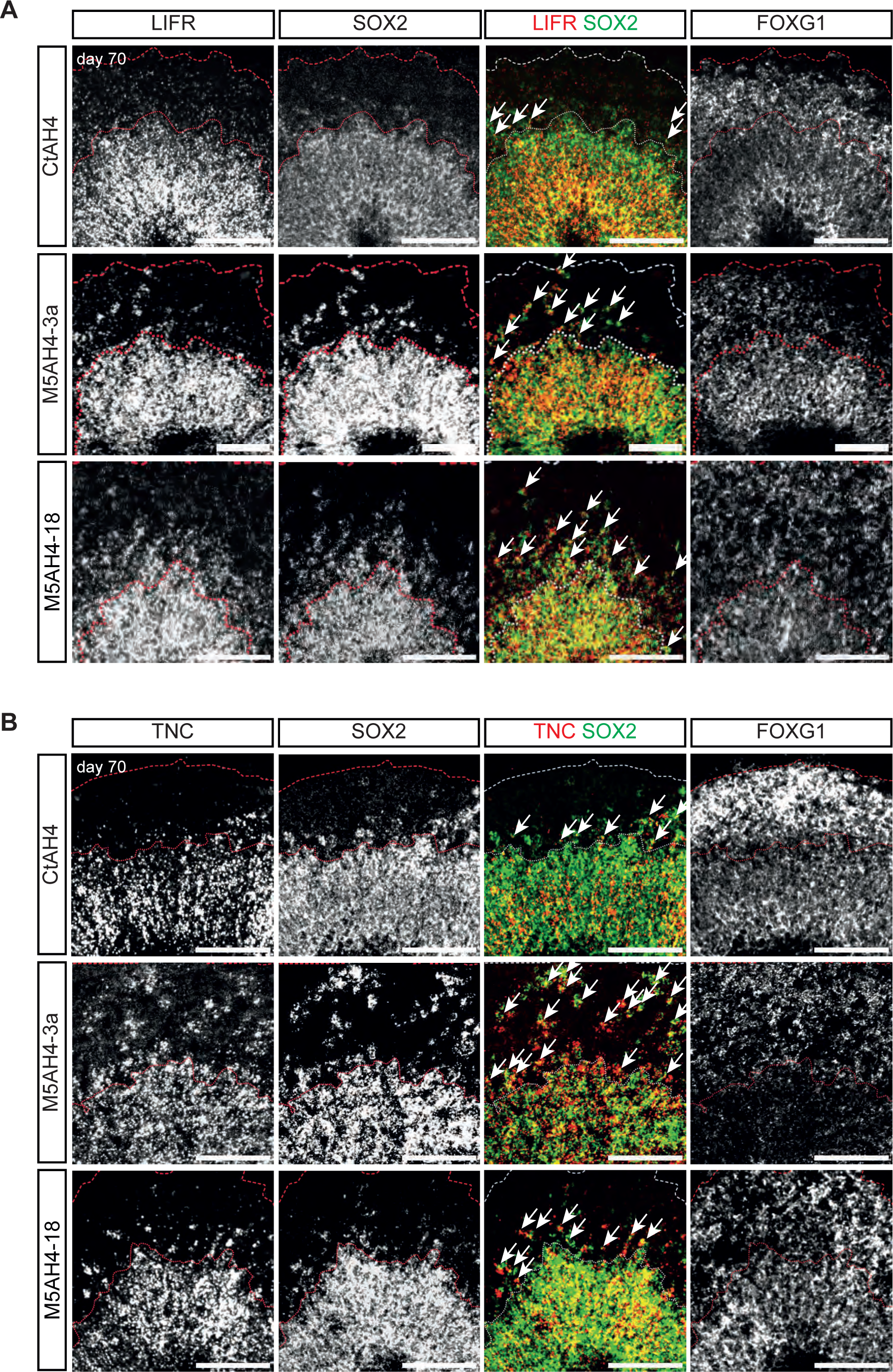
Delaminated SOX2+ cortical progenitors from the VZ in *ASPM*-mutated organoids display an oRG-like phenotype. (A and B) RNA-scope *in situ* hybridization showing that many displaced *SOX2*+ cells in *ASPM*-deficient organoids are co-expressing oRG-markers *LIFR* (A) or *TNC* (B) in VZ- like structures of organoids derived from CtAH4 and 5AH4-3a hiPSCs. All VZs have forebrain identity as shown by expression of *FOXG1*. Scale bars, 100 µm. See also Figure S6C.

Taken together, these data indicate that *ASPM*-mutated vRGC progenitors undergo a precocious delamination from the VZ and adopt an oRGC-like identity.

## Decreased amplification of vRGC and IPC in *ASPM*-mutant organoids

We next asked what could be the consequences of the precocious vRGC-to-oRGC conversion in the *ASPM*-mutant organoids. We first examined cell proliferation in *ASPM*- deficient organoids by quantifying PH3+ mitotic cells. We analyzed separately PH3+ cells touching the apical surface (“apical PH3+ cells”), and those located more basally within the VZ (“basal PH3+ cells) (Figures 6A–B and S7A–C). At day 35, the mitotic index was not different between both genotypes, regardless of the cells apical or basal position (Figures S7A–C). In contrast, at later stages, the *ASPM*-mutated organoids showed a strong decrease in the proportion of apical mitotic cells (Figures 6A–B), while the number of abventricular PH3+ cells remained unchanged (Figure S7B).

**Figure 6.**
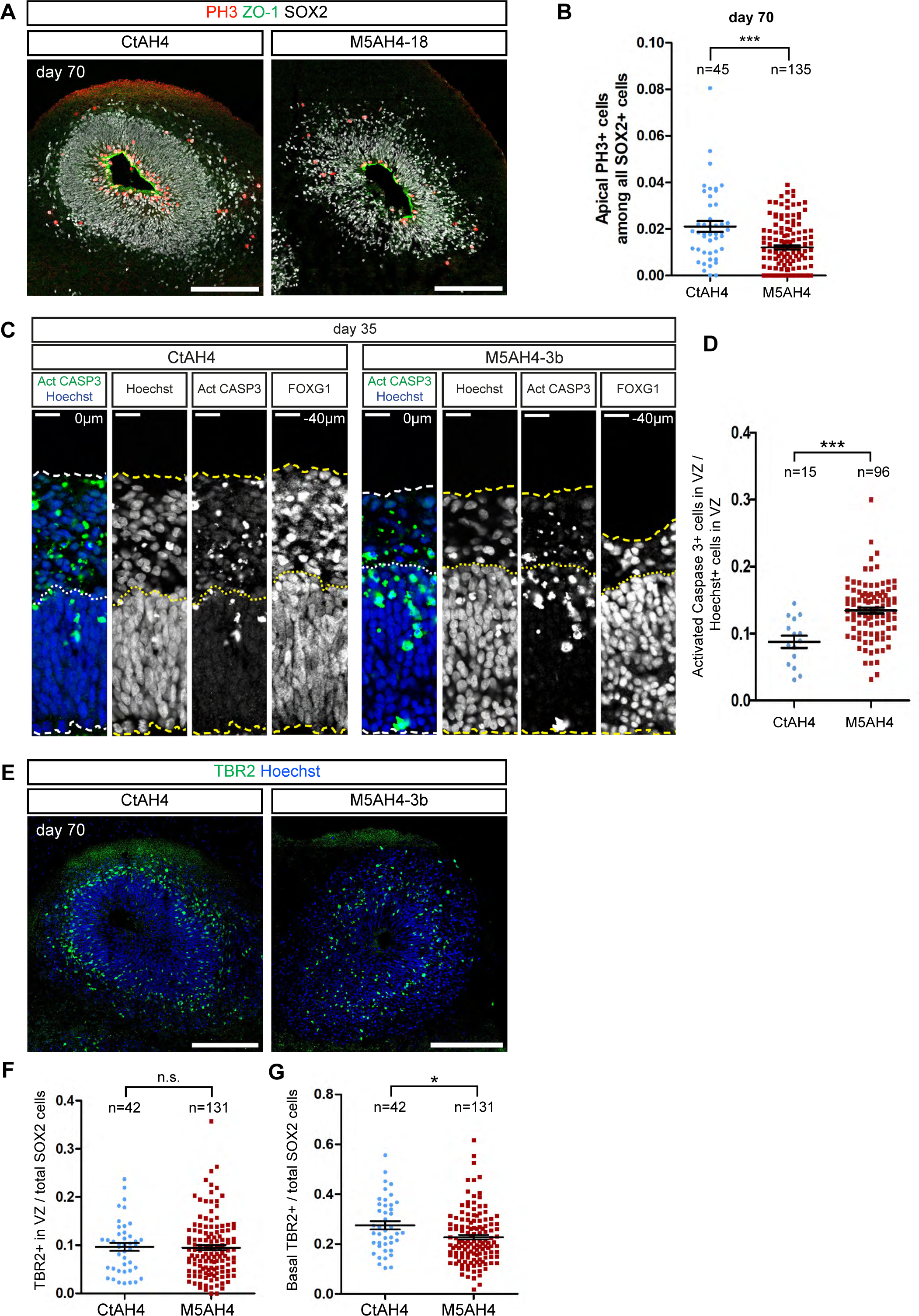
Late-stage impairment of vRG proliferation and IP generation, and early-stage increase in apoptosis of vRGs in *ASPM*-mutated organoids. (A) Representative confocal images of the distribution of PH3+ cortical progenitors (red) along the apical surface (ZO-1, green) in VZ-like structures of organoids derived from control and isogenic *ASPM*-deficient M5AH4-18. The VZ-like zone can be distinguished from the basally adjacent OSVZ-like zone by the density and coherence of SOX2+ nuclei (white). Although not different at day 35 (shown in Figure S7A), less apical PH3+ cells can be found by day 70 in the VZs of *ASPM*-mutated organoids compared with controls. (B) Scatter plots of the quantification of apical PH3+ progenitor cells (touching the apical surface) in cortical VZ-like structures at day 70, within a selected rectangle at the outer border of the organoid. The cortical identity was confirmed by FOXG1 staining on consecutive sections (not shown). One control and three isogenic *ASPM*-deficient AH4 cell lines from three experiments were analyzed with at least 12 VZ-like zones per experiment and cell line. Each dot represents one VZ. CtAH4: 18,925 SOX2+ in 7 organoids; M5AH4: 54,321 SOX2+ cells in 31 organoids counted. Shown above each genotype are the number of analyzed VZ-like structures for that line (n). Quantification of all abventricular PH3+ cells (not touching the apical surface, but located more basally in the VZ-like zone or residing in the OSVZ-like zone) and the total amount of PH3+ cells among all SOX2+ progenitor cells within a defined rectangle at day 35 and 70 are shown in Figures S7B and S7C. (C) Immunostaining showing an increased number of apoptotic (activated Caspase 3+, green) neural progenitors (Hoechst, blue) in *ASPM*-mutated M5AH4-3b organoid VZ-like structures with cortical identity as confirmed FOXG1+ on consecutive sections. The borders between VZ and OSVZ and the outside surface of the organoids are shown by dotted and dashed lines respectively. Similar observations shown for M5AH4-3a and M5AH4-18 in Figure S7D). (D) Scatter plot of the quantification of activated Caspase 3+ cells among all (Hoechst+) cells in selected 90° angles of VZs facing the outer border of the AH4 organoids showing more apoptotic cells in *ASPM*-mutated organoids at day 35. One control and three isogenic *ASPM*-deficient AH4 cell lines were analyzed. One dot represents one VZ. *P* value < 0.0001. CtAH4: 8 organoids from 4 experiments were analyzed, 3,393 Hoechst+ cells counted. M5AH4 lines: 40 organoids from 4 experiments were analyzed, 22,491 Hoechst+ cells counted. Mean ± SEM; Student’s *t*-test. (E) Representative immunostaining of TBR2+ intermediate progenitors in cortical VZ-like structures of isogenic AH4 organoids at day 70 showing a modest decrease in the number of intermediate progenitors in the OSVZ-like zone of *ASPM*-deficient organoids. (F and G) Scatter plots of the quantification of TBR2+ intermediate progenitor cells according to their position in cortical VZ-like structures of isogenic AH4 organoids within a selected 90° angle facing the outer border of the organoid at day 70. Quantification of the amount of TBR2+ cells in the VZ-like zone (F) and the amount of basally located TBR2+ cells located in the OSVZ-like zone (G) over the total amount of SOX2+ cells. One control and three isogenic *ASPM*-deficient AH4 cell lines from three experiments were analyzed with at least 12 VZ-like zones per experiment and cell line. CtAH4: 9 organoids, 9,683 SOX2+ cells; M5AH4 lines: 25 organoids, 13,132 SOX2+ cells. Mean ± SEM; Student’s *t*-test. Scale bars, 200 µm (A and E). Scale bars, 200µm (A and E), 20 µm (C).See also Figure S7.

We next analyzed apoptosis using immunostaining for activated Caspase 3, which revealed an increase in apoptotic cells of *ASPM*-mutated organoids at day 35, specifically within the VZ (Figures 6C–D and S7D–E). Finally, we examined TBR2+ IPC in isogenic control and *ASPM*-mutated AH4 organoids, which revealed a reduction of IPC in day 70 mutant organoids (Figures 6E–G, and S7F).

These data point to a depletion of VZ and SVZ progenitors following *ASPM*-deficiency, through a combination of decreased proliferation and increased apoptosis of vRG cells, as well as by the decrease in generation of IPCs.

## Discussion

*ASPM* mutations are the most frequent cause of MCPH^58^, but the mechanisms of this human disease have remained mostly unknown, especially since quantitatively and qualitatively distinct phenotypes were found in different animal models. Our findings point to a model whereby *ASPM* pathogenic mutations lead to precocious mitotic spindle randomization, and thereby to heterochronic generation of oRGC, followed by a depletion of vRGC and IPC. These data shed light on the mechanisms of MCPH and help explaining how specific mechanistic aspects of human corticogenesis may make our species particularly sensitive to *ASPM* disruption.

Using two different cortical organoid models, we found that mitotic spindle orientation changes dramatically during the early stages of corticogenesis. During the first 2-3 weeks, we observed a strictly horizontal mitotic spindle orientation in NEC and early vRGC, which then gradually evolved towards more randomized, non-horizontal orientations. These results are consistent with previous results obtained with human fetal cortex material, which also showed a gradual increase in the proportion of non-horizontal mitotic spindle pole in cortical progenitors, in correlation with increased generation of oRGC ^52^. Our data indicate that this early period of horizontal mitotic spindle orientation is critical for proper corticogenesis, as its disruption leads to precocious delamination of apical progenitors into oRGC-like cells.

On the other hand, in mouse cortical progenitors, the mitotic spindle remains more consistently horizontal throughout corticogenesis ^59,60^, but various mutant conditions associated with mitotic spindle misorientation can lead to delamination of vRGC into oRGC-like progenitors ^55,59^. In the ferret, *Aspm* disruption was shown to lead to early delamination of apical progenitors, but mitotic spindle misorientation was not explicitly reported ^39^. However, interestingly only late stages of corticogenesis were examined, and not those corresponding to very early stages including NEC, which were studied and revealed here.

Altogether these and our data thus suggest that cortical progenitors display species-specific temporal patterns of plane/mitotic spindle orientation, thereby leading to differential extent of delamination of apical progenitors towards basal compartments. This could explain how *ASPM* disruption, by influencing mitotic spindle orientation, can lead to distinct phenotypes depending on the species examined.

The precocious generation of oRGC-like progenitors in the *ASPM* mutants sheds further light on the species-specific sensitivity to *ASPM* dysfunction, but this observation also appears somewhat paradoxical: as oRGC are expected to drive cortical expansion, how can an increase in these cells be associated with MCPH? The most parsimonious explanation is that the *timing* of oRGC would be critical for proper cortical neurogenesis, by providing a match between specific types of RGC and specific niches found at different developmental stages in the apically located VZ and basally located OSVZ. Precociously displaced *ASPM*-mutant vRGC will develop in a heterotopic and heterochronic niche. They could therefore lack the potential to develop like normal oRGC and/or the niche could be unfit to sustain their proper lineage expansion and differentiation. Consistent with this hypothesis, studies in the ferret have identified a tight temporal window during which oRGC are generated, and which is required for proper neurogenesis ^61^. To address this important question it will be interesting to examine the impact of ASPM mutations on later stages of corticogenesis, using organoid models allowing long-term oRGC amplification, which remains challenging, or by examining *ASPM* mutant fetal brain pathology material, to determine what is the final outcome of the displaced oRGC in the mutant background. On the other hand, we found additional phenotypes in the vRGC, increased apoptosis and decreased proliferation, which lead to their precocious depletion. These phenotypes are likely related to the precocious displacement of the cells and deprivation from the apical niche, but they could also be linked to other effects of ASPM on cell mitosis and survival, which remains to be determined.

Another surprising observation of this study is that *ASPM* pathogenic mutations do not lead to complete loss of protein, whether in PSC or cortical progenitor cells. Rather our data indicate that truncated forms of ASPM are present in the mutant cells and are still properly localized at the mitotic spindle pole. *ASPM* mutations should therefore be considered as hypomorphic, or perhaps even neomorphic mutations, while fully null *ASPM* mutations in human cells remain to be generated. Interestingly, the various mutants generated display indistinguishable defects in corticogenesis, but the common feature between all *ASPM*-mutant cells in our study is the lack of expression of exon 18- encoded amino acids, corresponding to the IQ-domain. While the precise function of the IQ domain remains unknown, it was previously shown to be expanded further in larger brain-sized animals including primates and humans, and potentially the target of positive selection during human evolution^62^. While this hypothesis remains to be tested more directly, it is tempting to speculate that the IQ domain is somehow critical for the fine control of mitotic spindle orientation, which could be particularly critical in mammals with a large cerebral cortex.

Several other centrosome-associated MCPH genes (*CDK5RAP2*, *CPAP*, *STIL*, *WDR62*) have been studied in animal models and human PSC-based neural models ^58,63^. These studies reported diverse phenotypes including defects in neural stem cell mitosis, precocious differentiation, increased apoptosis, associated sometimes with mitotic spindle misorientation ^45,64–67^, but precocious generation of oRGC described here were not yet reported. Unlike *ASPM*, mutations in other MCPH genes can lead to severe phenotypes in the mouse, such as *CDK5RAP2* and *WDR62* ^35,68^. It will therefore be important to determine whether and how ASPM subserves functions that are distinct from other MCPH genes and that are required for normal cortical size in larger brain-sized animals, and thereby confers species-sensitivity to its disruption.

Although organoid systems provide unprecedented potential to monitor early events of human organ and tissue development *in vitro*, heterogeneity within and between organoids regarding cell type composition, the variable presence of non-cortical cells as well as culture condition-induced cell stress remain important caveats to consider in downstream analyses ^45,49,69^. Indeed the correct regional specification of neural progenitors was shown to be of critical importance for the modelling of *STIL*-gene linked MCPH phenotypes ^45^. Here we used a paradigm to generate human PSC-derived cortical tissue in a reproducible fashion from isogenic PSC-lines of various iPSC and ESC backgrounds. Importantly, we consistently focused our analyses on organoid regions displaying markers of cortical identity (FOXG1+/ EMX1+). Our approach illustrates how the use of consistent characterization of genomic and cell identity parameters appears to be an absolute prerequisite for the proper phenotypic studies of neural organoids, which should be refined further when studying even more complex phenotypes found in neurodevelopmental diseases.

In conclusion, *ASPM* pathogenic disruption results in early defects in the temporal transitions that orchestrate cortical neurogenesis, which eventually leads to the drastically decreased brain size found in the affected patients. Future work using pathological fetal material will be important to link more directly our *in vitro* approaches with the *in vivo* situation. On the other hand, further elucidation of the molecular mechanisms by which ASPM controls mitotic spindle orientation and apico-basal positioning of cortical progenitors will be required to fully understand how *ASPM* may have conferred our species distinct features of neurogenesis, as well as sensitivity to specific diseases.

## Acknowledgments

We thank members of the PV lab, IRIBHM, and CBD for helpful feedback and support, Kris Davie for help with scRNAseq and analyses, and Jean-Marie Vanderwinden of the Light Microscopy Facility (LiMiF) for support with imaging. We thank Catherine Verfaillie (KUL) for advice on iPSC reprogramming and Corinne Houart (King’s College) for helpful discussions. This work was funded by grants from the Belgian FNRS and FWO, the Belgian Queen Elizabeth Medical Foundation, the Interuniversity Attraction Poles Program (IUAP), the WELBIO Program of the Walloon Region, the Fondation de Spoelbergh, the AXA Research Fund, the Fondation ULB and the ERC (to P.V.), the Jean Van Damme fund of the Fonds Erasme (to M.A.), and the ERA-net E-Rare ‘EuroMicro’ (to M.A., S.P., and P.V.).

## Author Contributions

Conceptualization and methodology: AvB, RL, CL and PV; Investigation: AvB, RL, EE, DHT, AH, AB, LNR, MD, CL, MZ; Software: HvB, EE; Formal analysis: AvB, RL, EE, CL and PV; Key reagents: AvB, RL, MZ, IP, JD, MP, MA, SP, GM, KK, MZ, CL; Writing – original draft: AvB, RL, and PV; Writing – review and editing: AvB, RL, and PV; Funding: MA and PV; Resources: PV; Supervision: CL and PV

## Declaration of Interest

The authors declare no competing interests.

### STAR Methods

## RESOURCE AVAILABILITY

### Lead contact

Further information and requests for resources and reagents should be directed to Pierre Vanderhaeghen.

### Materials availability

All cell lines generated in this study will be shared by the lead contact upon request.

### Data and code availability

All data reported in this paper will be shared by the lead contact upon request. Any additional information required to reanalyze the data reported in this paper is available from the lead contact upon request.

## EXPERIMENTAL MODEL AND SUBJECT DETAILS

### Patients

Skin biopsies were obtained from patients diagnosed with mutations in *ASPM*, following a protocol approved by ULB and Erasme Hospital ethical committee. Table S2 provides an overview of the patients and their characteristics.

### iPSC line generation

Procedures of iPSC line generation and maintenance have been described in detail previously^41^. Briefly, Human dermal fibroblasts (HDF) from patients and control individuals (CtAH corresponds to HDF from a 32-year-old female donor and CtBJ to BJ1 fibroblasts derived from neonatal foreskin) were cultured in fibroblast medium (DMEM with 10 % FBS, 1% sodium pyruvate, 1 % penicillin/streptomycin (all from Thermo Fisher)). For their reprogramming, a retroviral infection system based on pMXs vectors carrying *c-MYC*, *KLF4*, *OCT4* and *SOX2* (Addgene) ^74,75^ was used. Each of the four viruses was produced by transfecting 6 x 10^6^ 293T cells with 75 μL FuGENE6 (Roche) or X- tremeGENE (Merck) and 100 μg DNA (pMX vector:gag-pol:vsv-g in a ratio of 4:3:1). Two days after transfection, the virus-containing supernatant was harvested on two consecutive days by filtration through a 0.45 μm cellulose acetate filter and 60-fold concentration using Amicon Ultra centrifugal filter units (Merck, 100 K). Fibroblasts were infected with fresh virus preparations one day after seeding them in a density of 100,000 cells/well of a 6-well plate with 5 μg/mL protamine sulfate (Merck). Four days after infection the cells were passaged on irradiated mouse embryonic fibroblasts (MEFs) and propagated in human ES medium (Knockout-DMEM containing 20% Knockout SR, 1% non-essential amino acids, 1% L-glutamine, 1% penicillin/streptomycin (all from Thermo Fisher), 0.1 mM 2-mercaptoethanol (Merck) and 10 ng/ml bFGF (Peprotech). iPSC colonies were isolated manually based on their morphology after 16–21 days. Chromosomal integrity was confirmed by karyotyping (Figure S1B).

### PSC culture

iPSC lines (AH4) and ESC lines (H9) were maintained on irradiated MEFs in hES medium using standard procedures. Cells were passaged enzymatically with Collagenase type IV/Dispase (1 mg/mL, each) every 4–5 days in the presence of 10 µM Y27632 (Abcam). To amplify the cells in feeder-free conditions they were transferred for one passage into dishes coated with hES-qualified Matrigel (Corning) with Essential 8 Medium (Thermo Fisher). For storage, hiPSC colonies were frozen in mFreSR medium (Stemcell Technologies) by putting the vials in an isopropanol freezing container at -80°C overnight before transferring them into liquid nitrogen. Pluripotency was confirmed by immunostaining (Figure S1C), embryoid body (Figure S1D) and teratoma formation (Figure S1E).

### Generation of homozygous insertion/deletion (INDEL) mutations in human *ASPM* by CRISPR/Cas9 editing

To generate INDEL mutations similar to the patient’s mutations, we targeted the human *ASPM* gene of control hiPSC line CtAH4 and hESC line CtH9 in exon 3 and 18 by CRISPR/Cas9-based gene editing (Figure 1A). For each of both exons a 20 nt single-guide RNA (crRNA) was designed based on the on-target and off-target score provided by Benchlinǵs bioinformatic output (https://benchling.com/, San Francisco, CA, USA) for the human *ASPM* gene^76,77^. Complementary pairs of oligonucleotides encoding the crRNA guide sequences were annealed and ligated separately into the BbsI site of the linearized plasmid eSpCas9(1.1) (Addgene; Plasmid #71814, ^78^), in which the sequence for TA2*-pac* gene was inserted downstream of the eSpCas9 gene to allow the selection of transfected cells with puromycin.

Next, the recombinant plasmid (1 µg) was nucleofected into 1–2 x 10^6^ CtAH4 and CtH9 cells using the Nucleofector™ 2b Device (Lonza) program B016 and Amaxa™ Human Stem Cell Kit 2 (Lonza; VPH-5052) according to the manufacturer’s protocol. The nucleofected cells were plated back in MEF-coated culture dishes and subjected to puromycin selection (0.5 µg/mL; InvivoGen) during 48h, starting 16h after nucleofection. After 10 more days, colonies were picked manually and expanded.

To ensure the successful introduction of homozygous mutations, the clones were analyzed by Sanger sequencing (Figure S1A) and ASPM immunostaining (Figure 1B and S1F). Pluripotency was confirmed by immunostaining (Figure S1C and D) and qRT-PCR (data not shown).

## METHOD DETAILS

### Generation of iPSC- and ESC-derived cortical organoids

Organoids with cortical identity were derived from entire stem cell (ESC/iPSC) colonies grown on irradiated MEFs, typically 5–7 days after passaging. On day 0, stem cell cultures were washed in sterile Dulbecco’s phosphate buffered saline (DPBS) and incubated at 37°C in a Collagenase type IV/Dispase solution (1 mg/mL each, Thermo Fisher) until colonies detached from the feeder layer (typically after 15’–20’). Detached colonies were gently collected and gravity sedimented in 5 mL human ES medium containing 10 µM Y27632, without FGF2, to neutralize the Collagenase type IV/Dispase by dilution. The colonies were washed once more in human ES medium as described above and subsequently in complete neural induction medium, consisting of DMEM/F12 + GlutaMAX supplemented with 20 % knock-out serum replacement, 1 % non-essential amino acids, 1 % penicillin-streptomycin and 100 µM 2-mercaptoethanol (all from Thermo Fisher), and further supplemented with 100 ng/mL Noggin (Bio-techne), 10 µM SB431542 (Stemcell Technologies), 2 µM (for iPSC) or 5 µM (for ESC) XAV939 (Stemcell Technologies), and 10 µM Y27632. Finally, colonies were resuspended in complete neural induction medium, plated in ultra-low attachment six-well plates (Greiner) and placed in stationary culture at 37°C and 5% CO2. Neural induction medium without Y27632 was renewed on day 1 and every other day onwards until day 7 (for ESC) or day 16 (for iPSC) included. At day 8 (for ESC) or day 17 (for iPSC), embryoid bodies were gently collected in 5 mL differentiation medium, consisting of DMEM/F12 + GlutaMAX as base medium supplemented with 1% non-essential amino acids, 1% penicillin-streptomycin, 100 µM 2-mercaptoethanol, 2.5 µg/mL insulin, 1% N2 and 2% B27 without vitamin A. After washing, embryoid bodies were resuspended in complete differentiation medium, plated in ultra-low attachment six-well plates (Greiner) and placed on a gyratory shaker (100 rpm) at 37°C and 5% CO2. Medium was renewed 3 times per week until the end of the experiment. From day 35 onwards, 1% vol/vol growth factor-reduced Matrigel (Corning) was added to the medium.

### PSC differentiation into 3DM cortical cultures

For neural differentiation in the 3DM system, feeder-based PSC colonies were passaged with Collagenase IV/Dispase (1 mg/mL, each) onto 6 well-plates covered with hES-qualified Matrigel and Essential 8 Medium to let them grow in feeder-free conditions. After 3–4 days, the colonies were washed with PBS, detached again with Collagenase IV/Dispase and washed twice with DPBS before embedding them in individual droplets of 50 µl of growth factor reduced (GFR) Matrigel (Corning). Each droplet was placed onto a glass coverslip. After gelling at 37°C for 10 min, the neural induction medium DDMB27 (consisting of DMEM/F12-GlutaMAX supplemented with 1x N2 supplement, 2 mM L-glutamine, 1x MEM-nonessential amino acids, 1 mM sodium pyruvate, 0.5 mg/mL BSA and 100 µm 2-Mercaptoethanol (all from Thermo Fisher))^41^ without any morphogens, but supplemented with BMP inhibitor Noggin (100 ng/ml, R&D Systems) ^44,79^ and Y27632 (10 µM, Abcam) was added. The culture medium was changed every other day by applying Noggin from day 0 to day 16 of the differentiation included.

After 21 days of differentiation in droplets of Matrigel, 3DM spheres were dissociated with Accutase and cells were seeded on coverslips coated with poly-L-Lysine/Laminin (Corning) in DDMB27 medium supplemented with Y27632 (10 µM). Five days after dissociation, half of the medium was exchanged against a mixture (1:1) of DDMB27:Neurobasal (Thermo Fisher) supplemented with B27 (without vitamin A, Thermo Fisher; 10 ml in 500 ml), 2 mM L-glutamine and 1% penicillin/streptomycin^41^.

### 2D PSC differentiation into cortical cells

The 2D differentiation protocol was slightly modified from what had been published earlier^41^. To remove the feeder cells from hES/hiPSC cultures, the colonies were passaged with Collagenase IV/Dispase (1 mg/mL, Thermo Fisher) into Essential 8 Medium (Thermo Fisher) on plates coated with hES-qualified Matrigel (Corning). After 3–4 days the colonies were dissociated using StemPro Accutase (Thermo Fisher), counted and plated at low confluency (10,000 cells/cm²) on Matrigel-coated dishes in Essential 8 Medium containing 10 µM ROCK inhibitor (Y27632). Two days later (day 0 of the differentiation), the medium was changed to DDMB27 with 100 ng/mL Noggin to start the neural induction. Until day 16, the medium was changed every other day, or— when indicated by the pH-indicator phenol red—every day. After day 16, the addition of Noggin to the DDMB27 was stopped and the medium was changed every other day. At day 25, the progenitors were washed with DPBS before incubating them for 5 min with StemPro Accutase at 37°C. Then, the cells were dissociated by gentle pipetting. The suspension was diluted with DDMB27 and centrifuged to remove the enzyme. To establish a stock of neural progenitors, the cell pellet from one 60 mm dish was resuspended in 2 mL mFreSR medium (Stemcell Technologies) to freeze two vials of the cells by putting them in an isopropanol freezing container at –80°C overnight before transferring them into liquid nitrogen.

To continue the culture, one vial of the frozen cells was thawed into two Matrigel-coated 60 mm dishes with DDMB27 Nb/B27 medium (one part of DDMB27 replenished with one part of Neurobasal supplemented with B27 (10 mL per 500 mL) and 2 mM L-glutamine. Every 2–3 days, the medium was partially changed by replacing half of the used medium.

### Immunofluorescence staining and quantifications

Organoids were fixed in 4% PFA overnight at 4°C after transferring them into a 15 mL-tube by using a wide bore pipet tip and washing them once with DPBS. The next day, they were transferred into 30% (vol/vol) sucrose/PBS for cryopreservation after three 5 min washes in PBS under gentle agitation. Following another night at 4°C, the organoids were embedded in Tissue freezing medium (Leica) on dry ice. The blocks were kept frozen at -80°C before cryosectioning at a thickness of 20 µm in RNase-free conditions. For immunostaining, the cryosections were washed during 10 min in PBS prior to a 1 h permeabilization/blocking step (with 3% BSA, 5% Horse serum, 0,3% Triton X-100 in PBS) and subsequent primary antibody incubation overnight at 4°C (in a solution of 3% BSA, 1% Horse serum, 0,1% Triton X-100). The next day, after three washes in PBS, cryosections were incubated at RT for 1 h in darkness with the secondary antibody and Hoechst 33258 (Merck) as a nuclear counterstain, washed again in PBS and finally mounted with a cover glass using Glycergel (Agilent Dako). Per slide, 500 µL of antibody solution were used.

Cells and 3DM spheres on coverslips were fixed with 4% paraformaldehyde (PFA) for 20 min or 1 h at RT, respectively. Cells intended for immunostaining with the N-terminal ASPM antibody SYR576 were fixed in ice-cold methanol for 5 min. For immunostaining of the 3DM spheres, the Matrigel droplets containing the fixed spheres were detached with a razor blade from the coverslips, embedded in blocks of 3% agarose and sectioned in 100 µm slices on a vibrating blade microtome (Leica) or mounted as whole droplets.

Immunofluorescence was performed as described above. A 1 h blocking step was followed by overnight incubation with the primary antibodies at 4°C. After three washing steps with PBS, incubation with the secondary antibodies was performed at RT for 1 h. All antibodies are listed in the tables below. Nuclei were visualized with Hoechst 33258. Glycergel (Agilent Dako) was used to mount the sections.

BrdU labeling was performed by incubating the 3DM spheres with 10 µM BrdU for 1 h before fixation with PFA for 4 h and absolute ethanol for 30 min at RT. Antigen retrieval was performed by incubating sections for 30 min in 2 N HCl at 37°C. The acid was neutralized by a 3% borate solution (pH 8.5) which was applied two times for 15 min.

### Custom-made N-terminal ASPM antibody

To enable the detection of the N-terminal domain of the ASPM protein upstream of the CRISPR/Cas9 editing, an ASPM antibody was generated by Eurogentec by using a 16 amino acid peptide (ac-SVSQKVDRVRSPLQAC-con) representing amino acids 180–195 of human ASPM (NP_060606.3). The antibody was raised in rat against the KHL- conjugated peptide. After confirming its affinity (Eurogentec; Peptide: 1712973, Rat SYR576) through ELISA, the antibody was affinity matrix purified.

### Quantification of cells in organoids

All quantifications were performed in organoid regions with dorsal forebrain identity as characterized on consecutive sections by positive immunoreactivity for FOXG1 and EMX1. For quantifications, VZ-like structures facing the outer borders of the organoids were imaged by confocal microscopy.

In case of the quantification of SOX2+, TBR2+ and activated Caspase 3+ cells, an angle of 90° was drawn in FIJI (ImageJ) by placing the shortest distance to the pial surface in its middle and the vertex of the angle at the apical surface (see illustration Figure 4B). Within this angle, the border between the cell-dense VZ-like region and the basal OSVZ- like region was defined by using the density and coherence of the SOX2+ nuclei as a criterion. SOX2+ and TBR2+ cells were counted in each compartment and related to the total number of SOX2+ cells. Activated Caspase-3+ cells were counted within the densely packed VZ-like zone and related to the total number of Hoechst+ nuclei present within the VZ-like zone.

In case of the quantification of PH3+ cells, a rectangle instead of an angle was used for analysis, as the angle vertex converging to the apical surface would miss most of the mitotic PH3+ cells that are inherently located at the apical surface. The rectangle was placed at the half of the VZ-like structure facing the outer border of the organoid. Again the border between the VZ-like and the OSVZ-like compartment was delineated according to the SOX2+ cell density and coherence.

All experiments were done in triplicate by running control and *ASPM*-mutated cells in parallel. For the isogenic AH4 cell lines, one control and three *ASPM*-mutated cell lines were analyzed. The quantification results were plotted in Prism (GraphPad).

### RNAscope

To detect mRNA expression of *LIFR* and *TCN* in organoids, *in situ* hybridization was performed on 4% PFA-fixed cryosections (prepared as described above for immunofluorescence) using the RNAscope Multiplex Fluorescent Reagent Kit v2 (Advanced Cell Diagnostics) according to the user manual. Sections were post-fixed with 4% PFA for 15 min at RT and subsequently dehydrated in 50%, 70% and 100% ethanol for 3 min each. After letting the sections dry at RT, they were incubated for 10 min with 5–8 droplets of H2O2. For antigen retrieval, the slides were treated for 5 min with Target Retrieval Reagent. After 3 min in 100% ethanol, slides were dried again and a barrier was drawn around the section with a hydrophobic barrier pen. Next, the sections were pretreated with Protease III for 10 min in a HybEZ oven at 40°C to improve antigen accessibility. After washing with PBS, the sections were hybridized with a probe mix containing FOXG1-C3, SOX2-C2/-C1, and LIFR-C2 or TNC-C1 probes for 2 h in the oven at 40°C. The slides were rinsed twice with 1x Wash Buffer at RT, before incubating them sequentially with Multiplex FLv2 Amp1/Amp2/Amp3 solution at 40°C for signal amplification. To label the amplified hybridizations with fluorophores, the slides were incubated sequentially with Multiplex FLv2 HRP-C1, -C2 and -C3 for 15 min at 40°C, with Opal 570, Opal 520 or Opal 690 diluted (1:1500) in 1x Plus Amplification Diluent for 30 min at 40°C and with Multiplex FLv2 HRP blocker for 15 min at 40°C. Nuclei were labeled with DAPI (1:2000). Coverslips were mounted on the slides with ProLong Gold Antifade Mountant (Thermo Fisher). All steps until the probe hybridization were performed under RNAse-free conditions.

### Embryoid body formation

hiPSC colonies were detached with collagenase IV and transferred into Costar Ultra low attachment 6-well plates (Corning) where they were cultured in suspension in hES medium without the addition of bFGF. After 10 days the embryoid bodies were dissociated with 0.05% trypsin/0.5 mM EDTA, plated on gelatin-coated coverslips and grown for 10 more days.

### Teratoma formation assay

iPSC from feeder-free cultures were enzymatically dissociated to single cells with StemPro Accutase (Thermo Fisher), washed and resuspended in 50 µl of Matrigel. At least 1 x 10^6^ cells were injected subcutaneously into the dorsal flank of immunocompromised NOD-SCID mice. Tumors were isolated 8–12 weeks after injection and fixed in 4% paraformaldehyde overnight. After embedding in paraffin and sectioning, the teratomas were analyzed based on hematoxylin and eosin staining. All mouse experiments were performed with the approval of the Université Libre de Bruxelles Committee for animal welfare.

### Karyotyping

iPSC colonies were incubated overnight in feeder-free conditions with 0.05 µg/ml KaryoMAX Colcemid solution (Thermo Fisher). The next morning the cells were washed with PBS and detached with trypsin/EDTA. After dissociation, the cells were centrifuged, resuspended and incubated in 2 ml pre-warmed 0.075 M hypotonic KCl solution for 30 min at 37°C. The cells were fixed by dropwise addition of 0.5 ml freshly prepared ice-cold methanol:glacial acetic acid (3:1), centrifuged and resuspended in 1 ml fixative. The last step was repeated twice. A cytogenetic lab (IPG, Gosselies, Belgium) spread and analyzed the cells after Giemsa staining.

### Spindle orientation

Usual two-dimensional analysis of the mitotic spindle orientation is reported to be error-prone and can lead to incorrect results, since it neglects the spindle orientation in z-direction and the unevenness of the apical surface ^80,81^. For this reason, we did a 3D analysis by reconstructing confocal z-stacks with Imaris 3D Image analysis software (Bitplane) and subsequent computational calculation.

On organoid sections, all analyses of spindle orientation were performed in regions with forebrain identity as characterized on consecutive sections by positive immunoreactivity for FOXG1.

Sections of cortical organoids and 3DM spheres were immunostained with nuclear mitotic apparatus protein (NuMA) to label centrosomes, with phospho-Vimentin, a marker for dividing progenitor cells in the developing neocortex ^82^, with PH3 to distinguish mitotic phases more easily, and counterstained with Hoechst to label chromatin. Phospho-Vimentin staining did not only mark the cytoplasm of mitotic cells, but also revealed the contour of the apical surface. This allowed us to calculate a plane in the apical surface underneath the dividing cell, in order to determine the orientation of the cell in relation to this plane. Therefore, confocal images with z-stacks of 500 nm were taken and three-dimensionally reconstructed using Imaris 3D Image analysis software (Bitplane). First, the dividing cell and the apical surface were reconstructed separately by manual retracing of the pVimentin staining on consecutive z-stacks of the dividing cell and at the apical surface. The intersection of the two defined volumes of the dividing cell and the apical surface was calculated allowing the export of the resulting coordinates (x, y and z) into Excel. Second, the center of mass of each chromatin half of the dividing cell was calculated through automatic recognition by the Imaris software and the coordinates of the two resulting points were exported into Excel. To analyze the spindle orientation, a plane with the intersecting points of the dividing cell and the apical surface was calculated by applying an ordinary least squares regression analysis to the surface coordinates. At least 20 intersection points were used per cell to calculate the plane. Next, the cleavage angle orientation α of the mitotic spindle (Figure 4B) was derived from the angle β between the plane’s normal vector and the vector between the two chromatin halves and is defined by α = 90° – β, meaning that values close to 0 degrees represent a horizontal and values close to 90 degrees a vertical division in relation to the apical surface. To perform this analysis a program was written in Haskell that uses the Excel files as input and returns the spindle orientation α (see Program code S1). To verify the fit of the plane, the program also generates a 3D visualization showing the coordinates obtained from Imaris, the calculated the plane and the vector between the two chromatin halves. Ten cells were analyzed per cell line and experiment. In case of the isogenic AH4 cell lines, one control and three *ASPM*-mutated cell lines were analyzed in the organoid and the 3DM system; in case of the patient-derived hiPSC lines, four control cell lines and seven *ASPM*-mutated cell lines were analyzed in the 3DM system.

### Confocal and widefield microscopy

Confocal images were generated with a Zeiss LSM780 confocal system fitted on an Observer Z1 inverted microscope equipped driven by ZEN 2012 software (Zeiss) and equipped with 5x/0.16, 10x/0.3, 20x/0.8 and 40x/1.1W objectives and a Chameleon Vision II 690–1064 nm multiphoton laser (Coherent Europe).

Widefield images were generated on a Zeiss Axio with 5x and 10x objectives. Images were processed using Fiji/ImageJ software.

### Organoid dissociation and scRNA-seq library preparation

Organoids derived in parallel from the iPS lines CtAH4 and M5AH4-3a at day 36 (Amount of organoids per cell line: CtAH4: 8, M5AH4-3a: 4), day 56 (CtAH4: 5, M5AH4-3a: 5) and day 70 (CtAH4: 5, M5AH4-3a: 5) were processed for single cell RNA-sequencing (scRNA-seq) as follows. Organoids were collected using a wide bore pipette tip at the respective time points, washed in HBSS, snapped into two pieces using a sterile blade in order to initially remove debris from the organoid cores ^83^, washed twice in HBSS and incubated during 30 min at 37°C in pre-warmed, oxygenated (95% O2, 5% CO2) Papain/DNase I (Worthington Biochemical Corporation) on a gyratory shaker (100 rpm). A single cell suspension was prepared by gentle trituration and centrifugation (300 *g*, 5 min). Debris was further removed by subsequent gradient centrifugation on Ovomucoid protease inhibitor/DNase I (70 *g*, 6 min), cells were resuspended in PBS containing 0.04% bovine serum albumin on ice, and finally filtered to remove doublets prior to cell counting and viability assessment.

### Single cell RNA sequencing

Library preparations for the single cell RNA-seq were performed using 10X Genomics Chromium Single Cell 3’ Kit, v3.1 Next GEM chemistry (10X Genomics, Pleasanton, CA, USA). The cell count and the viability of the samples were accessed using LUNA dual fluorescence cell counter (Logos Biosystems) and only samples with viability >80% were processed for downstream 10X processing with a targeted cell recovery of 10,000 cells for each of the samples. After cell count and QC, the samples were immediately loaded onto the Chromium Controller. Single cell RNA-seq libraries were prepared using manufacturers recommendations (Single cell 3’ reagent kits v3.1 user guide; CG000204 Rev D), and at the different check points the library quality was accessed using Qubit (Thermo Fisher) and Bioanalyzer (Agilent).

10X libraries were either sequenced on Novaseq 6000 (Illumina) or MGISEQ-2000 (BGI) sequencing platforms. Illumina sequencing was done as per 10X genomics recommendation: paired end read of 28 bps (read 1), 91 bps (read 2) and 10 bps (index 1). MGISEQ-2000 sequencing was done at the MGI Hong Kong SAR sequencing facility. Before being able to sequence on the MGI platforms, the 10X libraries need to undergo a conversion step using MGIEasy Universal DNA Library Prep kit. Briefly, the final 10X sequencing libraries were circularized using the splint-ligation step and the circularized libraries were converted to single-stranded DNA copies. DNA nanoballs were prepared from the circularized ssDNA using Rolling Circle Amplification (RCA). DNB libraries generated were then flown through the patterned flow cell of the MGISEQ- 2000RS High-Throughput Sequencing kit. For a targeted sequencing saturation of 50– 60%, sequencing was performed at a depth of 30,000–60,000 reads per cell. A custom sequencing recipe of paired end read of 28 bps (read 1), 100 bps (read 2) and 10 bps (index 1) was used for sequencing.

### Pre-processing of single cell data

10x Genomics single cell samples were demultiplexed using the 10x Genomics Cell Ranger v6.1.2. Cellranger *count* command was used for mapping and generation of gene expression matrices with GRCh38 Human genome was used as reference genome (provided by 10X Genomics).

### Analysis and integration of single cell RNA organoid data

Filtered gene expression matrices were processed using the Scanpy (1.8.2) package ^84^. Cells with less than 1000 genes or more than 15% of counts present in mitochondrial genes were excluded, counts were log-normalized and counts per cell were scaled to a total of 10,000. Highly variable genes were selected with *pp.highly_variable_genes* using default settings and principal component analysis (PCA) was performed with the selected genes. PCs were adjusted between organoid samples using Harmony with *external.pp.harmony_integrate* ^85^. The top 40 PCs were selected and used to compute a neighborhood graph. UMAP was computed using default Scanpy parameters. Unsupervised clustering was performed using the Louvain algorithm (resolution=2). Differentially expressed genes cell types were obtained by via *tl.rank_genes_groups* using Wilcoxon method for each cluster. Scaled expression of cell type marker genes (obtained from ^45^) were visualized using matrixplot after scaling the gene expression matrices with *pp.scale*.

The raw and normalized data of developing human cortex ^49^ was downloaded from UCSC Cell Browser (https://cells.ucsc.edu; dataset ID: ‘organoidreportcard/primary10X’). The normalized expression matrix was subset to select only CS22 stage cells. Interneurons, outlier cells, cells with less than 500 genes and more than 15% mitochondrial counts were removed from the dataset. Organoid and *in vivo* expression matrices were concatenated after log normalization and highly variable genes were selected for PCA. PCs were adjusted between *in vivo* and *in vitro* batches using Harmony with *external.pp.harmony_integrate*. Corrected PCA embeddings were then used to form the neighborhood graph and compute UMAP as above.

## QUANTIFICATION AND STATISTICAL ANALYSIS

### Statistical analysis

For statistical analysis Student’s unpaired t-test was performed to calculate *P*-values and the results were indicated as * for p<0.05; ** for p<0.01; *** for p<0.001 and n.s. for not significant. Results are shown as mean ± standard error (SEM).

## Supplementary item titles

Program code S1 (related to Figure 4): Program for the analysis of the cleavage angle of mitotic cells

## Supplementary figure titles and legends

**Figure S1.**
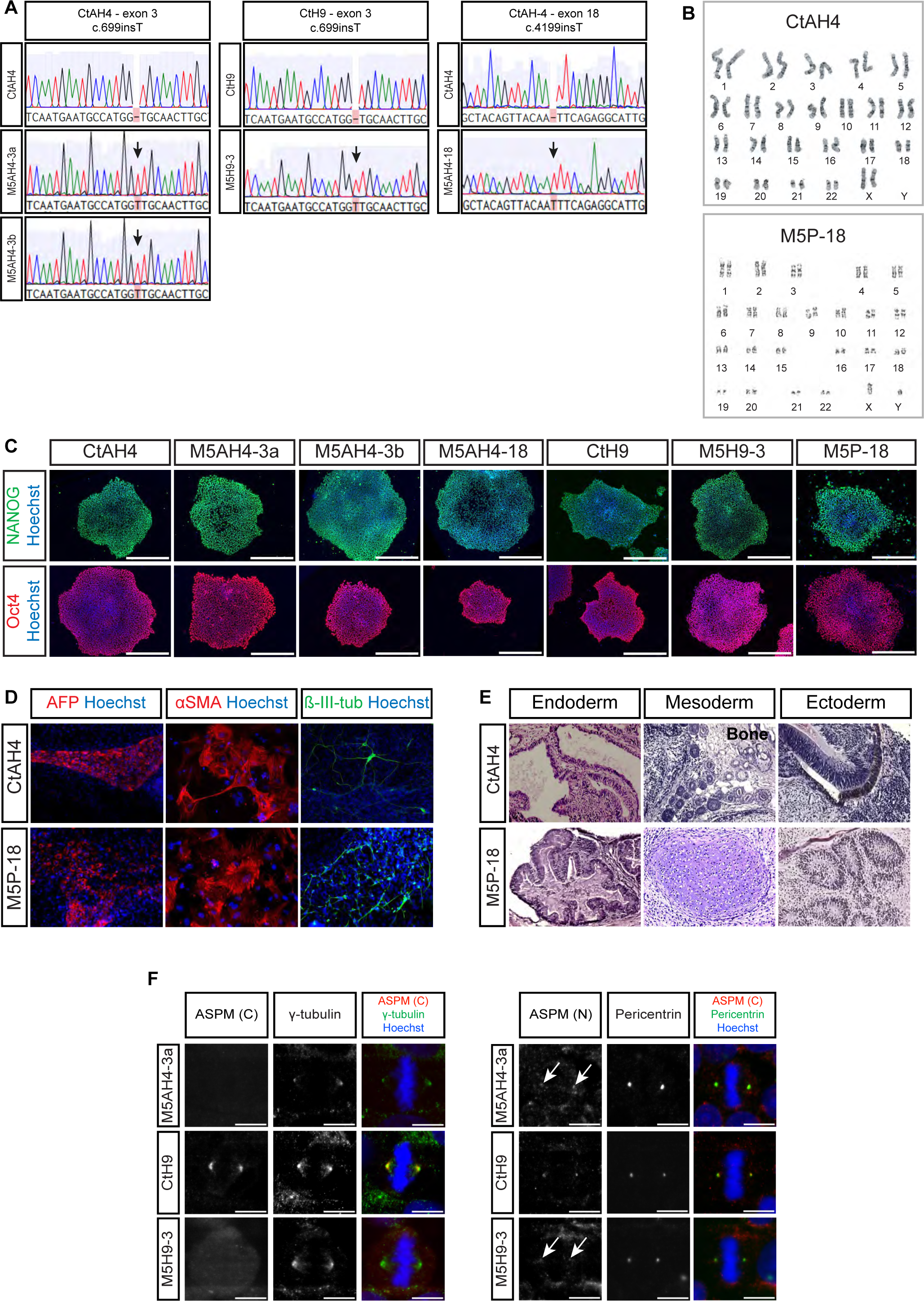
(related to Figure 1). Characterization of *ASPM* mutations in isogenic hiPSC and hESC lines, pluripotency validation of PSC lines and ASPM protein expression in isogenic PSC lines and patient-derived iPSCs. (A) Sequencing results of hiPSC line AH4 and hESC line H9 before and after introducing *ASPM* mutations by CRISPR/Cas9 gene editing showing the unchanged control (top row) and the isogenic cell lines mutated by insertion of a T in exon 3 or 18 (arrow, bottom rows). The generated mutations are homozygous. For an overview of all mutations in the isogenic hiPSC/hESC lines and MCPH patients of this study see also Table S1 and S2 respectively. (B) Karyotypes revealing chromosomal integrity shown for control hiPSC and patient M5P-18 analyzed by G-banding. (C) Representative immunofluorescence staining of pluripotency markers NANOG and OCT4 on all isogenic PSC lines and patient line M5P-18. Nuclei are counterstained with Hoechst. (D) Immunofluorescence analysis following differentiation of control hiPSCs and patient line M5P-18 into embryoid bodies shows generation of endoderm (α-fetoprotein, AFP), mesoderm (α-smooth muscle actin, α-SMA) and ectoderm (β-III-tubulin). Nuclei were counterstained with Hoechst. Embryoid bodies were cultured in suspension for ten days before dissociation and passaging on gelatin-coated coverslips. Images were taken at day 20. (E) Hematoxylin and eosin staining of sections from hiPSC- and patient line M5P-18- derived teratomas revealing differentiated tissues from all germ layers: endoderm is indicated by the presence of glandular structures, mesoderm by the presence of cartilage (not noted in the image), bone, adipose tissue or muscle and ectoderm by the presence of (pigmented) neuroepithelium and neural rosettes. Teratomas were analyzed 8–12 weeks after injection of iPSCs into NOD-SCID mice. (F) Related to Figure 1B, subcellular localization of ASPM protein in mitotic hiPSCs with a homozygous mutation in exon 3 (M5AH4-3a), control hESCs (CtH9) and isogenic hESCs with a homozygous mutation in exon 3 (M5H9-3). Scale bars, 500µm (C) and 10µm (F).

**Figure S2.**
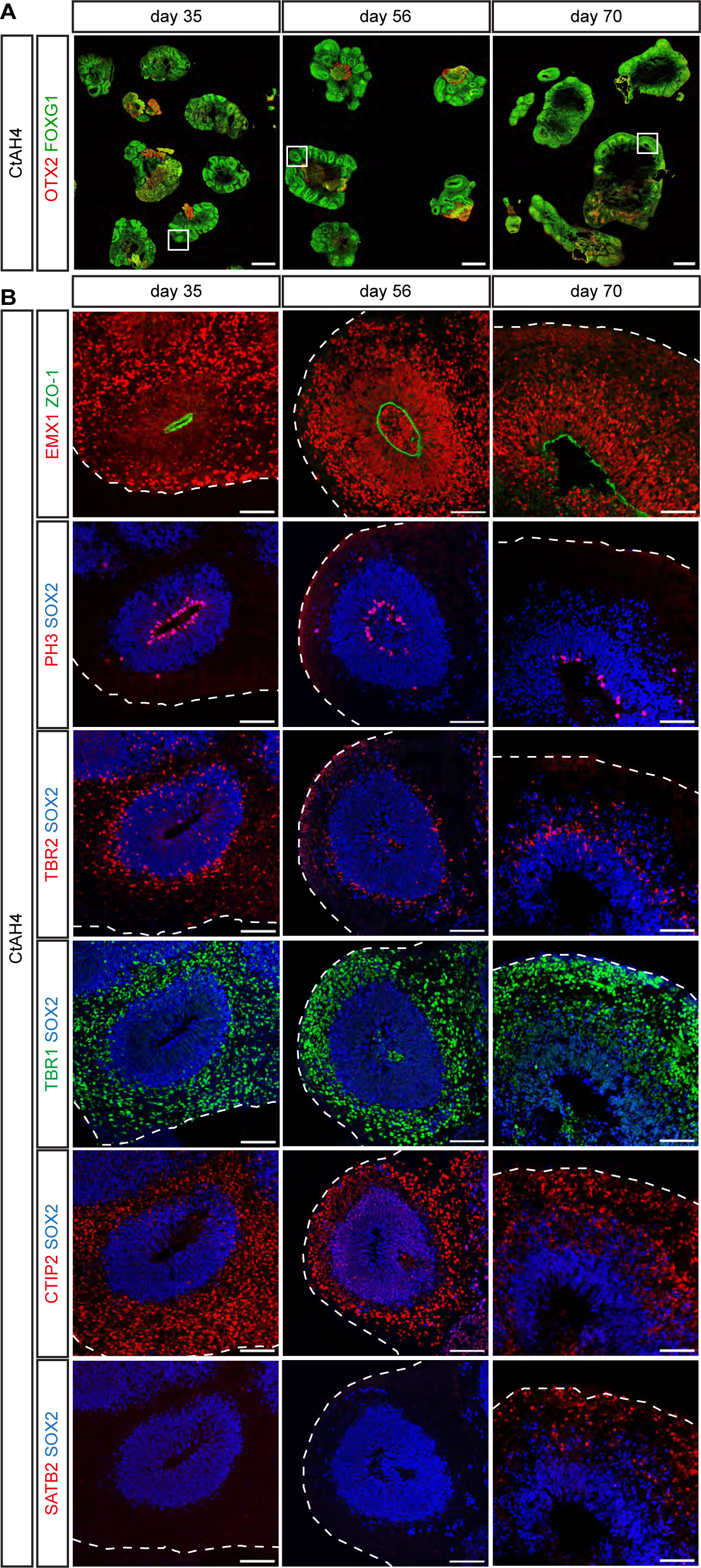
(related to Figure 2). Detailed characterization of marker expression in cortical organoids from control hiPSCs. (A) Overview images of the generation of organoids with cortical identity from control CtAH4 hiPSCs showing sustained FOXG1 expression over time at day 35, 56, and 70. (B) Boxed VZs from the overview of organoids in (A) are shown in detail for different markers on consecutive sections characterizing the apico-basal organization of the cell-dense VZ (ZO-1; PH3; SOX2), its cortical fate (EMX1), and the successful generation of IPCs (TBR2+), deep-layer neurons (TBR1+; CTIP2+) and upper-layer neurons (SATB2+) over time (day 35, 56, and 70). The outer borders of the organoids are indicated by dashed lines. Scale bars, 1 mm (A), 100 µm (B).

**Figure S3.**
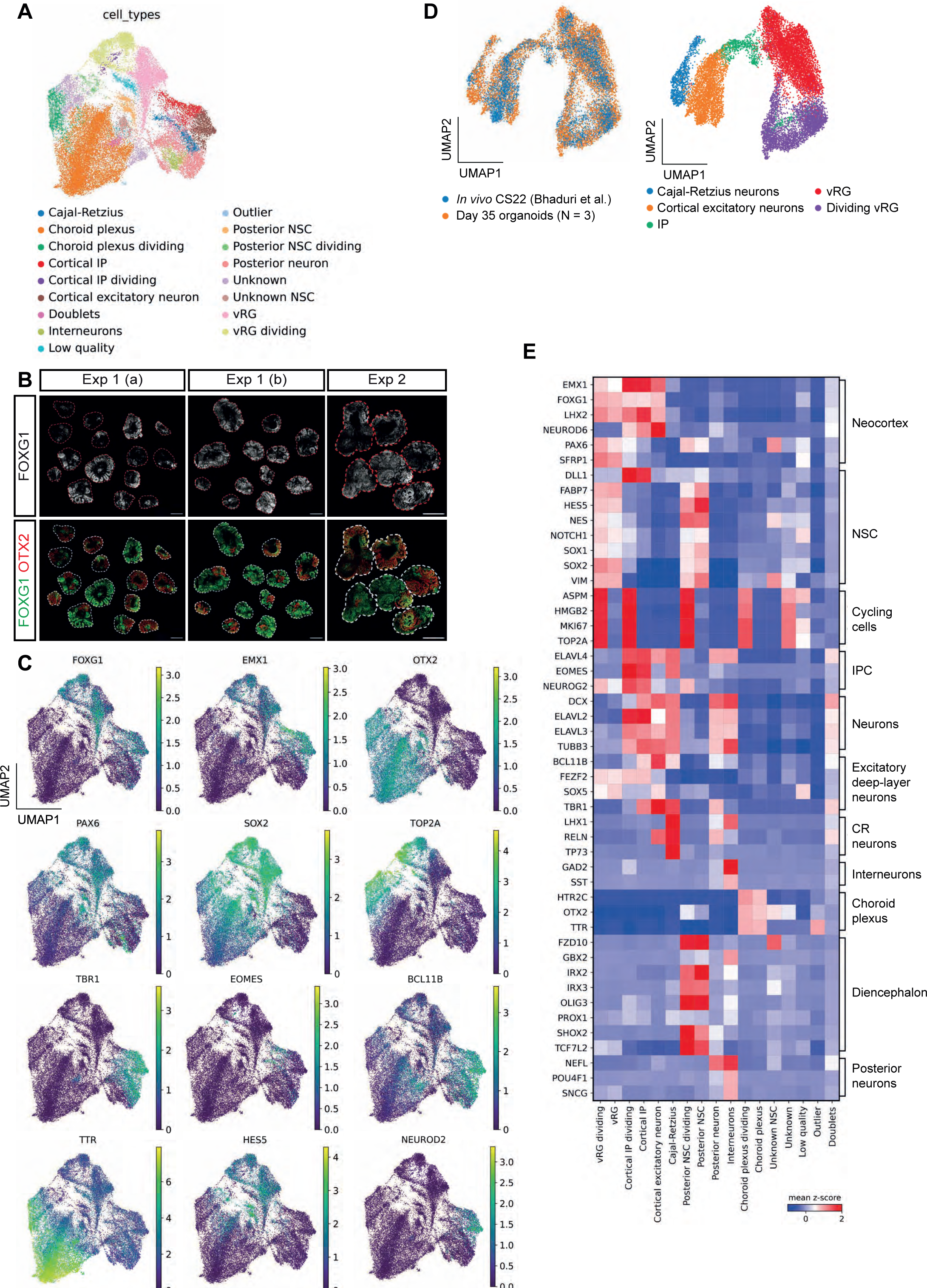
(related to Figure 2). Transcriptomic characterization and validation of cortical organoids by scRNAseq. (A) Uniform manifold approximation and projection (UMAP) derived from CtAH4 organoids from three biological replicates showing unsupervised clustering using the Louvain method. (B) Example overview immunofluorescence images depicting intra- and inter-experimental variability in the generation of FOXG1+ and OTX2+ regional specifications. Shown are two sets of organoids within the same experiment (exp 1a and 1b), and a third set of organoids from a different experiment (exp 2). Top row: single channel FOXG1 in grey, bottom row: merged FOXG1/OTX2 in green and red respectively, showing heterogeneous generation of OTX2+ regions. Outer borders of the organoids are indicated by dashed lines. (C) Single marker profiles on the same UMAP embedding as in (A). (D) Merged transcriptomic data from cortical cells of day 35 CtAH4 organoids derived from (A) and CS22 *in vivo* human cortex ^49^ show overlap of cortical cell types. (E) Matrix plot of scaled expression of marker genes used in Rosebrock et al. ^45^.

**Figure S4.**
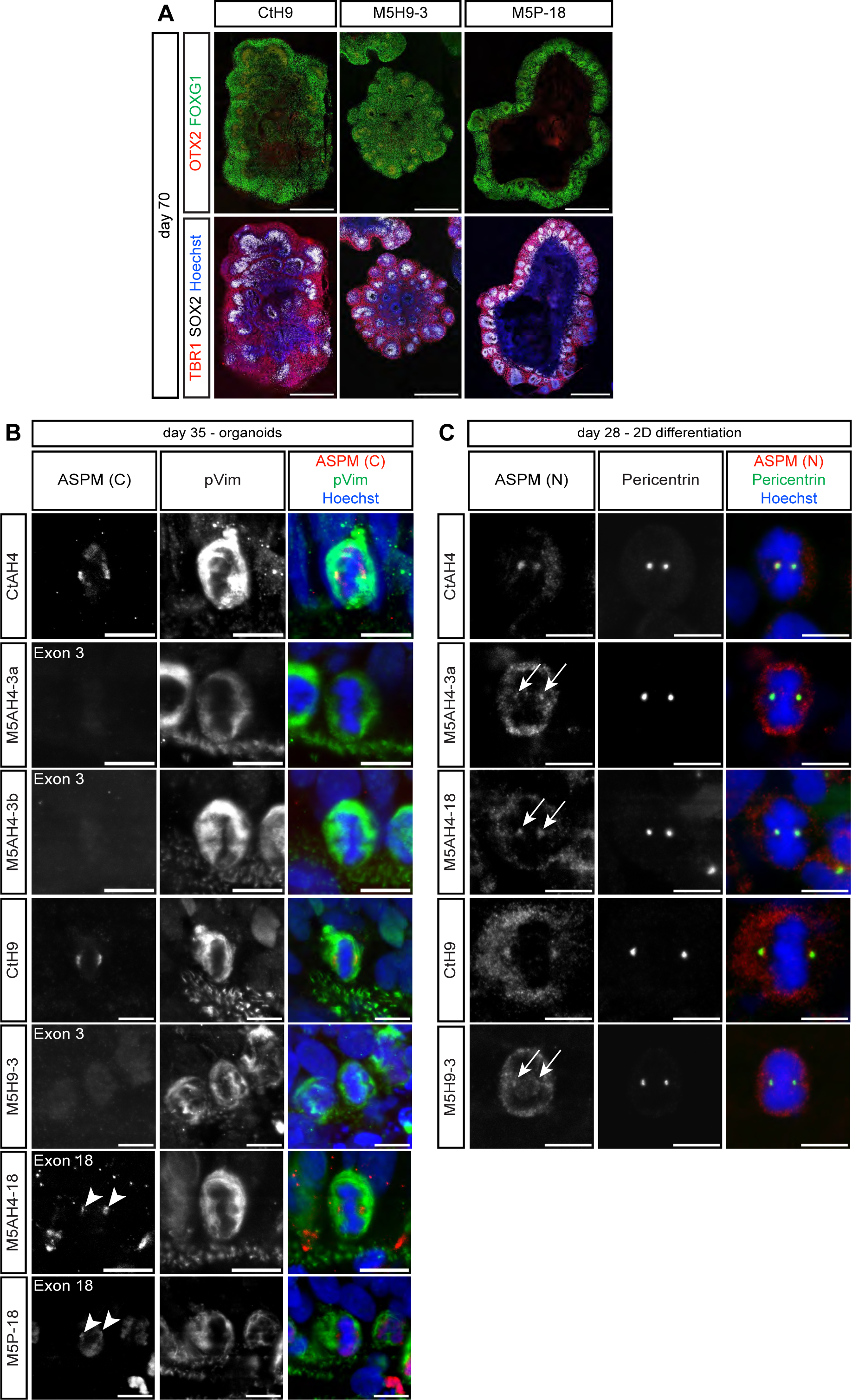
(related to Figure 1 and 2). Characterization of marker expression in cortical organoids from control hESCs, *ASPM* exon 3-mutated hESCS and *ASPM* exon 18-mutated patient iPSCs and subcellular localization of ASPM protein mitotic neural progenitors derived from isogenic hiPSC-/ hESC- and patient-derived cell lines. (A) Immunofluorescence staining of single organoids of the isogenic hESC lines (CtH9 and M5H9-3) and the patient-derived hiPSC line M5P-18 showing relevant cortical cytoarchitecture and marker expression at day 70 of differentiation. Shown are FOXG1+ VZ-like zones facing the outer borders of the organoids (top row) and TBR1+ deep-layer cortical neurons, arranged basally around SOX2+ progenitors within the VZ-like zones. As shown for the other isogenic hiPSC lines in Figure 2A and 2B cytoarchitecture and marker expression are similar across genotypes. (B) Localization of ASPM protein in mitotic neural progenitors in organoids derived from isogenic control and *ASPM*-mutated AH4 hiPSCs and H9 hESCs and from the patient hiPSCs M5P-18 (mutated in exon 18) at day 35 revealed by the C-terminal ASPM antibody. As seen in hiPSCs/hESCs, the control cells show a strong immunoreactivity at the spindle poles, while labeling is less intense in exon 18- and absent in exon 3-mutated cell lines. pVimentin (green) reveals vRG cells in M phase. Merged images additionally show Hoechst staining in blue. (C) Localization of ASPM protein in mitotic neural progenitors in 2D differentiations of isogenic control and *ASPM*-mutated AH4 hiPSCs and H9 hESCs at day 28 revealed by the N-terminal ASPM antibody (merged image: red). Although immunoreactivity for ASPM is detected at the spindle poles in all conditions, the signal is less intense in the *ASPM*- mutated cells. Merged images show centrosomal pericentrin in green and Hoechst in blue. Scale bars, 1000 µm (A), 10 µm (B and C).

**Figure S5.**
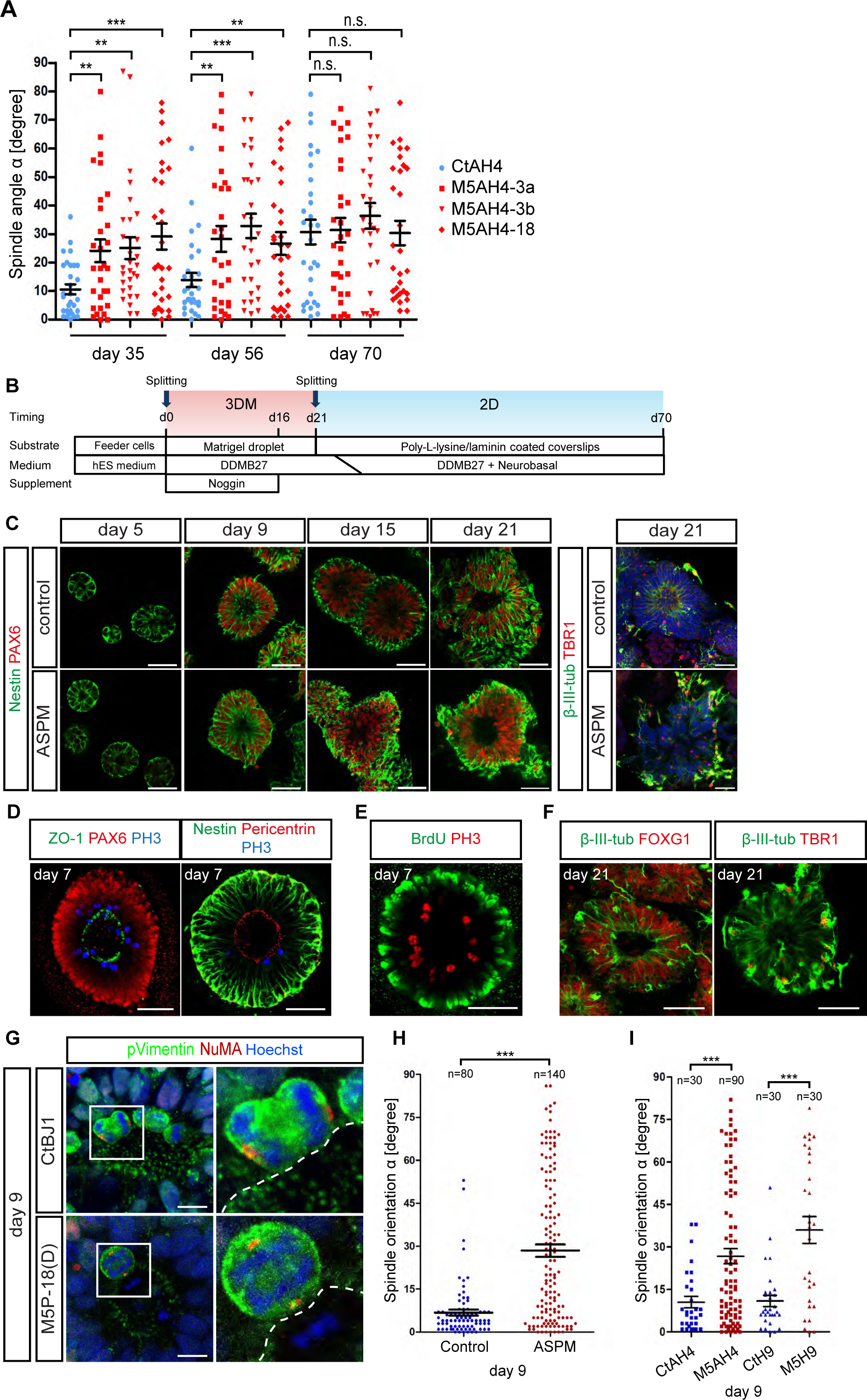
(related to Figure 3). Premature alteration of spindle orientation in *ASPM*- deficient neural progenitors of cortical organoids and early-stage 3DM corticospheres. (A) Scatter plot showing the quantification of cleavage angle orientation in isogenic control and *ASPM*-deficient AH4 progenitor cells analyzed by 3D reconstruction at day 35, 56 and 70 of organoid differentiation, displayed per single M5AH4 cell line. These results are shown as pooled for all M5AH4 isogenic *ASPM*-mutant cell lines in Figure 3C. This highlights that all of the isogenic *ASPM*-mutated cell lines show significantly more oblique/vertical cleavage angles at day 35 and 56 of the organoid differentiations compared to the control, while by day 70 of differentiation the control cells have reached similar levels of spindle angle randomization (n = 30 for all cell lines and time points). Mean ± SEM; Student’s *t*-test. (B) Schematic presentation of the procedure for generating cortical progenitors and neurons with the 3DM differentiation system starting from iPSCs embedded in GFR Matrigel. (C) Left: generation of Nestin-positive 3DM spheres which start to express the cortical progenitor marker PAX6 between day 5 and day 9 indicating an identity of early forebrain progenitors. Right: immunostaining of early born β-III-tubulin-positive neurons in control and *ASPM*-mutated 3DM spheres at day 21. Some of the early-born neurons from both conditions co-express the glutamatergic neocortical marker TBR1. (D) Apico-basal polarity in the 3DM spheres at day 7 revealed by expression of ZO-1 and Pericentrin at the apical surface. (E) BrdU labeling for 1 h before fixation reveals NE cells in the 3DM spheres at day 7 undergoing INM as indicated by BrdU-positive cells residing at the basal surface and mitotic PH3-positive cells at the apical surface. (F) The first TBR1-positive neurons, observed around day 21, appear in the outer borders of the 3DM spheres and indicate cortical identity (FOXG1+/TBR1+). (G) Representative images of mitotic neural progenitors in anaphase in control (CtBJ1) and *ASPM*-mutated (M5P19) 3DM spheres at day 9. Shown are overview immunofluorescence stainings for pVimentin and NuMA on the left with higher magnification of the boxed area in the right panel. Dashed lines indicate the apical surface. While the majority of mitotic cells in the control condition divides in parallel to the apical surface, the number of randomized (“oblique” and “vertical”) cleavage angles is increased in *ASPM*-deficient cells. (H) Scatter plot showing the quantification of cleavage angle orientation in control and *ASPM*-deficient patient-derived NE cells analyzed by 3D reconstruction at day 9 of 3DM differentiation. Most of the cortical progenitors of the control group divide horizontally (mean α = 6.81 ± 1.05 SEM), while *ASPM*-mutated neural progenitors show a premature increase in oblique/vertical cleavage angels (mean α = 28.46 ± 2.15 SEM). Analysis included four CtAH and CtBJ control cell lines (see Methods, iPSC line generation) and seven *ASPM*-mutated patient-derived cell lines from two experiments (see Table S2). (I) Scatter plot showing the quantification of cleavage angle orientation in isogenic control and *ASPM*-deficient NE cells from AH4 hiPSCs and H9 hESCs analyzed by 3D reconstruction at day 9 of 3DM differentiation. Most of the cortical progenitors of the control groups divide horizontally (CtAH4: mean α = 10.50 ± 2.00 SEM; CtH9: mean α = 10.90 ± 2.00 SEM), while *ASPM*-mutated neural progenitors show more randomized (“oblique” or “vertical”) cleavage angles (M5AH4: mean α = 26.72 ± 2.64 SEM; M5H9: mean α = 35.97 ± 4.74 SEM). One control (CtAH4) and three isogenic *ASPM*-mutated AH4 cell lines, and one control (CtH9) and one isogenic H9 cell line from three experiments, respectively (see Table S1). Mean ± SEM; Student’s *t*-test. Scale bars, 50µm (C) and 10µm (D)

**Figure S6.**
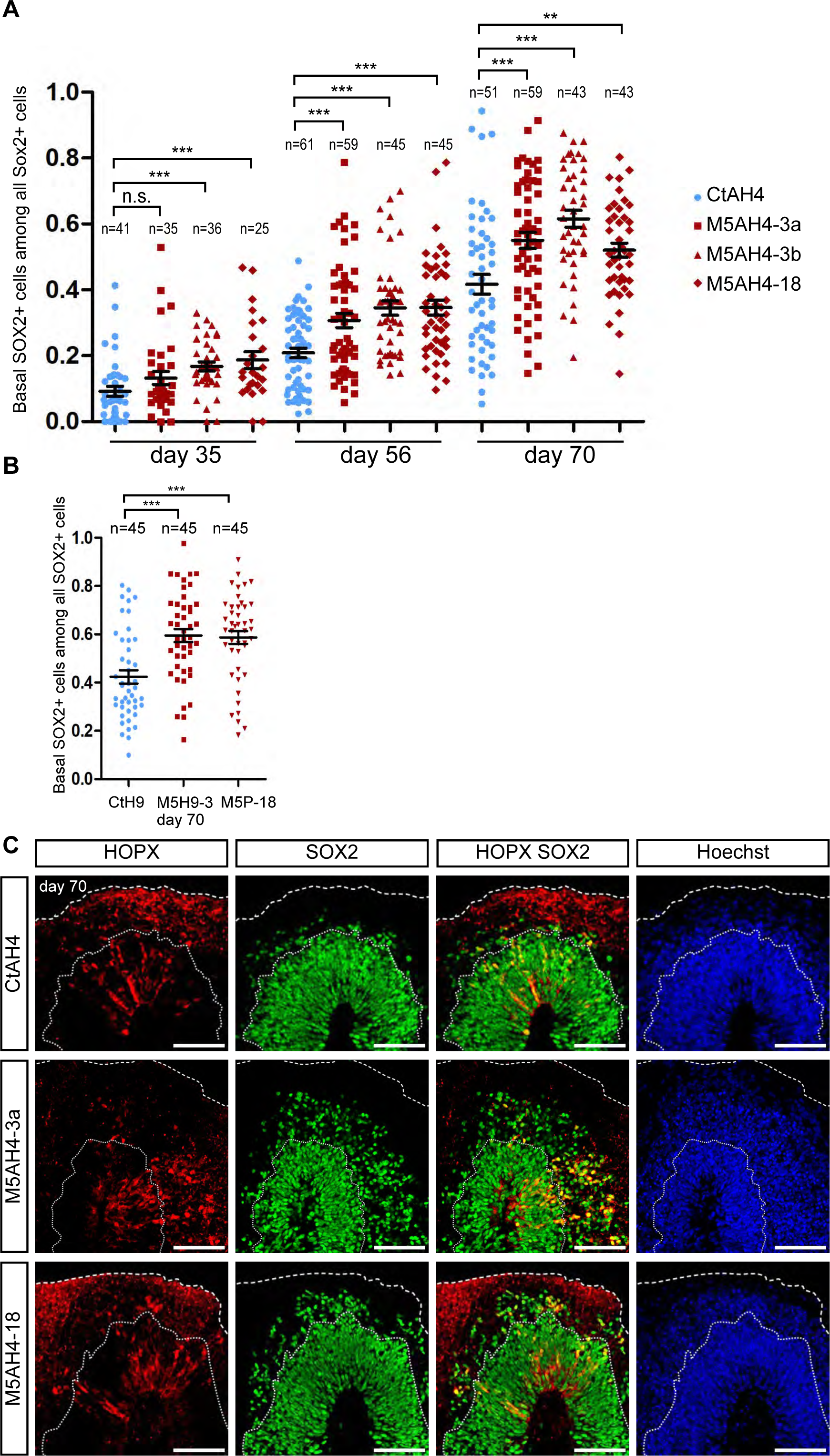
(related to Figures 4 and 5). SOX2+ cortical progenitors precociously delaminate from the VZ-like zones in *ASPM*-mutated organoids and display an oRG-like phenotype. (A) Scatter plot showing the quantification of basal SOX2+ cells delaminated from the VZ-like zone among the total amount of SOX2+ cells in isogenic AH4 organoids at day 35, 56 and 70, displayed for the CtAH4 control line and each *ASPM*-mutated M5AH4 line separately. Shown above each cell line are the number of analyzed VZ-like structures for that line (n). These results are shown as pooled for all M5AH4 isogenic *ASPM*-mutant cell lines in Figure 4C. (B) Same analysis as in (A) for the isogenic ESC cell lines (CtH9 and M5H9-3) and the patient-derived iPSC line with an exon 18 *ASPM*-mutation (M5P-18), at day 70. For all three cell lines 45 VZs from three experiments were analyzed. Both *P* values < 0.0001; CtH9: 19 organoids, 5,512 SOX2+ cells; M5H9-3: 23 organoids, 5,191 SOX2+ cells; M5P-18: 23 organoids, 8,331 cells. (C) Immunostaining showing an increased number of displaced SOX2+ cells in *ASPM*- mutated organoids co-expressing the oRG marker HOPX.

**Figure S7.**
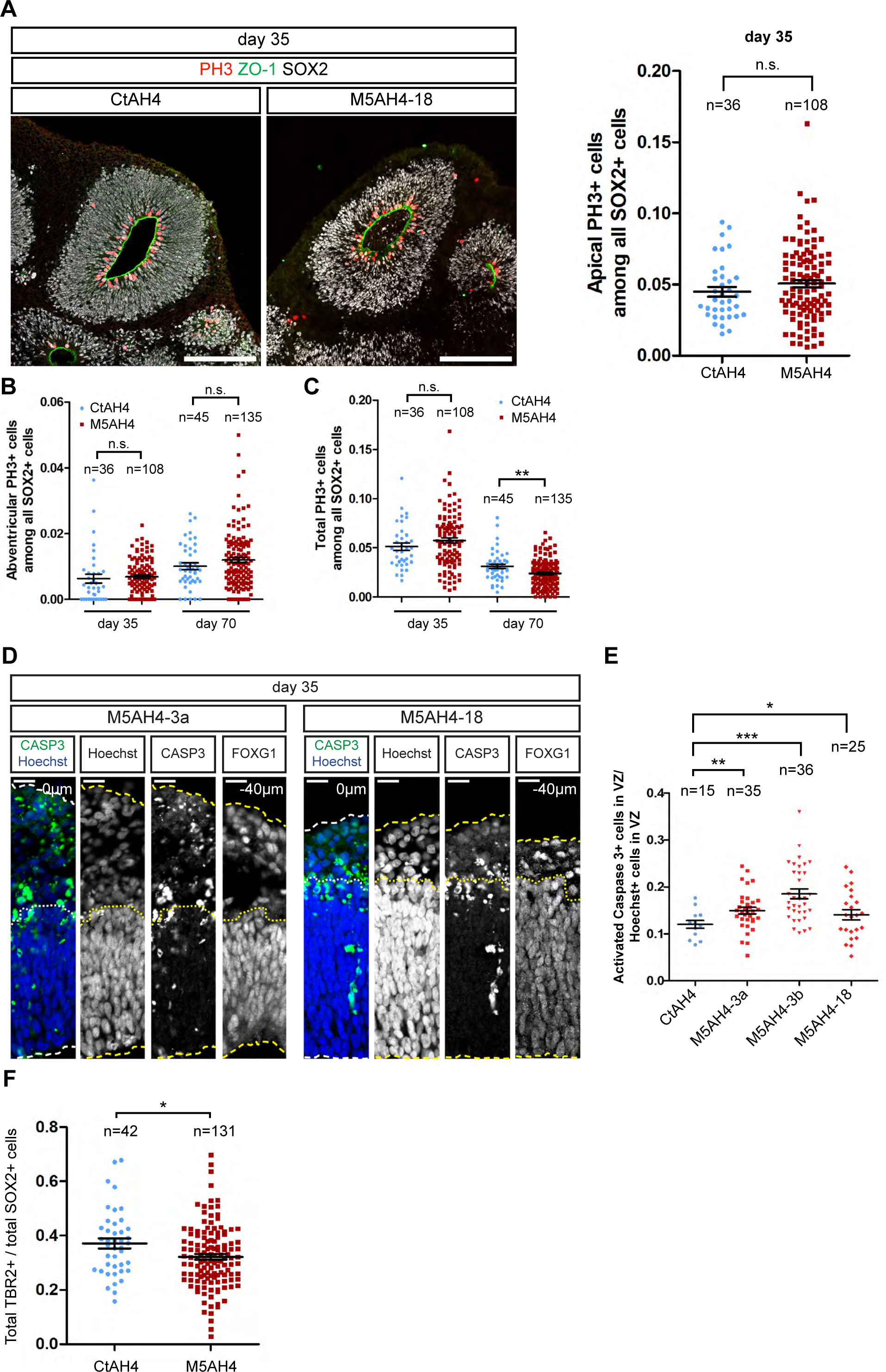
(related to Figure 6). Late-stage impairment of vRG proliferation and IP generation, and early-stage increase in apoptosis of vRGs in *ASPM*-mutated organoids. (A) Left: representative confocal images of the distribution of PH3+ cortical progenitors (red) along the apical surface (ZO-1, green) in VZ-like structures of day 35 organoids derived from control and isogenic *ASPM*-deficient M5AH4-18. The VZ-like zone can be distinguished from the basally adjacent OSVZ-like zone by the density and coherence of SOX2+ nuclei (white). Right: scatter plot of the quantification of apical PH3+ progenitor cells (touching the apical surface) in cortical VZ-like structures at day 35, within a selected rectangle at the outer border of the organoid. The cortical identity was confirmed by FOXG1 staining on consecutive sections (not shown). One control and three isogenic *ASPM*-deficient AH4 cell lines from three experiments were analyzed with at least 12 VZ-like zones per experiment and cell line. Each dot represents one VZ. CtAH4: 12,476 SOX2+ in 7 organoids; M5AH4: 39,819 SOX2+ cells in 31 organoids counted. Shown above each genotype are the number of analyzed VZ-like structures for that line (n). (B and C) Quantification of all abventricular PH3+ cells (not touching the apical surface, but located more basally in the VZ-like zone or residing in the OSVZ-like zone) and the total amount of PH3+ cells among all SOX2+ progenitor cells within a defined rectangle at day 35 and 70. One control and three isogenic *ASPM*-mutated AH4 cell lines from three experiments were analyzed with at least 12 VZ-like zones per experiment and cell line. Each dot represents one VZ. Day 35: CtAH4: 12,476 SOX2+ in 7 organoids; M5AH4: 39,819 SOX2+ cells in 31 organoids counted. Day 70: CtAH4: 18,925 SOX2+ in 7 organoids; M5AH4: 54,321 SOX2+ cells in 31 organoids counted. Shown above each genotype are the number of analyzed VZ-like structures for that line (n). (C) Immunostaining showing an increased number of apoptotic (activated Caspase 3+, green) neural progenitors (Hoechst, blue) in *ASPM*-mutated M5AH4-3a and M5AH4-18- derived organoid VZ-like structures with cortical identity as confirmed FOXG1+ on consecutive sections, similar to the observations shown in Figure 6C. The borders between VZ and OSVZ and the outside surface of the organoids are shown by dotted and dashed lines respectively. (D) Scatter plot of the quantification of activated Caspase 3+ cells among all (Hoechst+) cells in selected 90° angles of VZs facing the outer border of the AH4 organoids showing more apoptotic cells in *ASPM*-mutated organoids at day 35, displayed per single M5AH4 cell line. One control and three isogenic *ASPM*-deficient AH4 cell lines were analyzed. One dot represents one VZ. *P* value < 0.0001. CtAH4: 8 organoids from 4 experiments were analyzed, 3,393 Hoechst+ cells counted. M5AH4 lines: 40 organoids from 4 experiments were analyzed, 22,491 Hoechst+ cells counted. Mean ± SEM; Student’s *t*-test. These results are shown as pooled for all M5AH4 isogenic *ASPM*-mutant cell lines in Figure 6D. (E) Scatter plots of the quantification of the total amount of TBR2+ intermediate progenitor cells over the total amount of SOX2+ cells in isogenic AH4 organoids within a selected 90° angle facing the outer border of the organoid at day 70. Quantification of the total amount of TBR2+ cells over the total amount of SOX2+ cells. One control and three isogenic *ASPM*-mutated AH4 cell lines from three experiments were analyzed with at least 12 VZ-like zones per experiment and cell line. CtAH4: 9 organoids, 9,683 SOX2+ cells; M5AH4 lines: 25 organoids, 13,132 SOX2+ cells. Shown above each genotype are the number of analyzed VZ-like structures for that line (n). Mean ± SEM; Student’s *t*-test. Scale bars, 200 µm (A), 20 µm (D).

Table S1: Isogenic cell lines.

Table S1: Isogenic cell lines
| Cell line name | M5AH4-3a | M5AH4-3b | M5H9-3 | M5AH4-18 |
| --- | --- | --- | --- | --- |
| **Mutations in ASPM** | Homozygous | Homozygous | Homozygous | Homozygous |
| **Exon** | 3 | 3 | 3 | 18 |
| **cDNA** | c.699insT | c.699insT | c.699insT | c.4199insT |
| **Protein** | p.Gln233GlnfsX22 | p.Gln233GlnfsX22 | p.Gln233GlnfsX22 | p.Ile1400IlefsX17 |

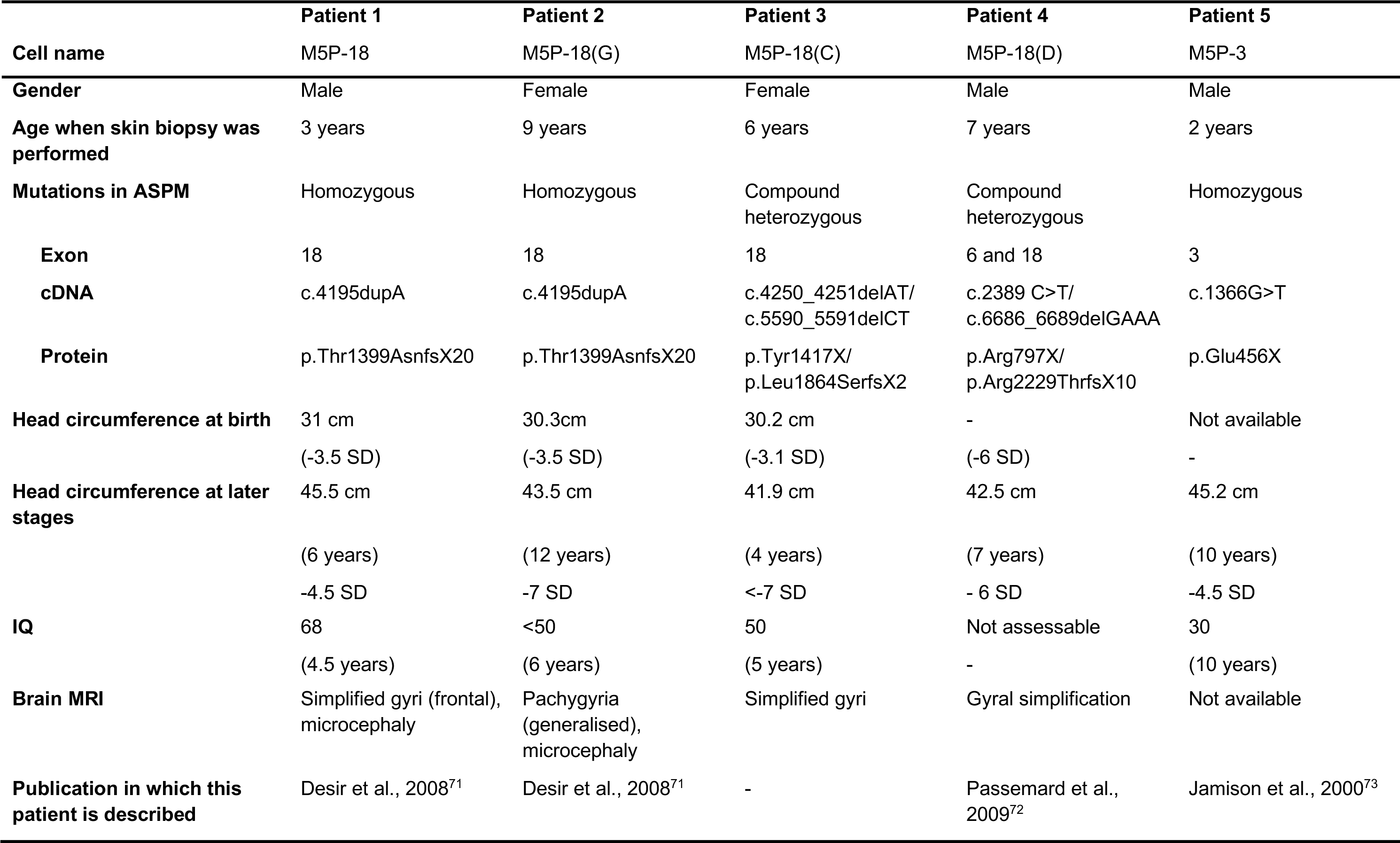
Table S2: Characterization of MCPH patient.

## Notes

### Competing Interest Statement

The authors have declared no competing interest.

